# Rewiring of 3D chromatin topology orchestrates transcriptional reprogramming in muscle fiber-type specification and transformation

**DOI:** 10.1101/2025.03.16.643571

**Authors:** Baohua Tan, Ting Gu, Linjun Hong, Liyao Xiao, Jiajin Wu, Geyan Lu, Shanshan Wang, Langqing Liu, Enqin Zheng, Gengyuan Cai, Zicong Li, Zhenfang Wu

## Abstract

The composition of muscle fibers, characterized by distinct contractile and metabolic properties, significantly influences meat quality and glucose homeostasis. However, the mechanisms by which three-dimensional (3D) genome topology integrates with epigenetic states to regulate muscle fiber specification and transformation remain poorly understood. Here, we present an integrative analysis of the transcriptome, epigenome, and 3D genome architecture in the slow-twitch glycolytic extensor digitorum longus (EDL) and fast-twitch oxidative soleus (SOL) muscles. Global remodeling of enhancer-promoter (E-P) interactions emerged as a central driver of transcriptional reprogramming associated with muscle contraction and glucose metabolism. We identified tissue-specific super-enhancers (SEs) that regulate muscle fiber-type specification through cooperation of chromatin looping and transcription factors such as KLF5. Notably, the SE-driven activation of *STARD7* facilitated the transformation of glycolytic fibers into oxidative fibers by mitigating reactive oxygen species levels and suppressing ERK MAPK signaling. This study elucidates the principles of 3D genome organization in the epigenetic regulation of muscle fiber specification and transformation, providing a foundation for novel therapeutic strategies targeting metabolic disorders and enhancing meat quality.

Schematic overview of the study.

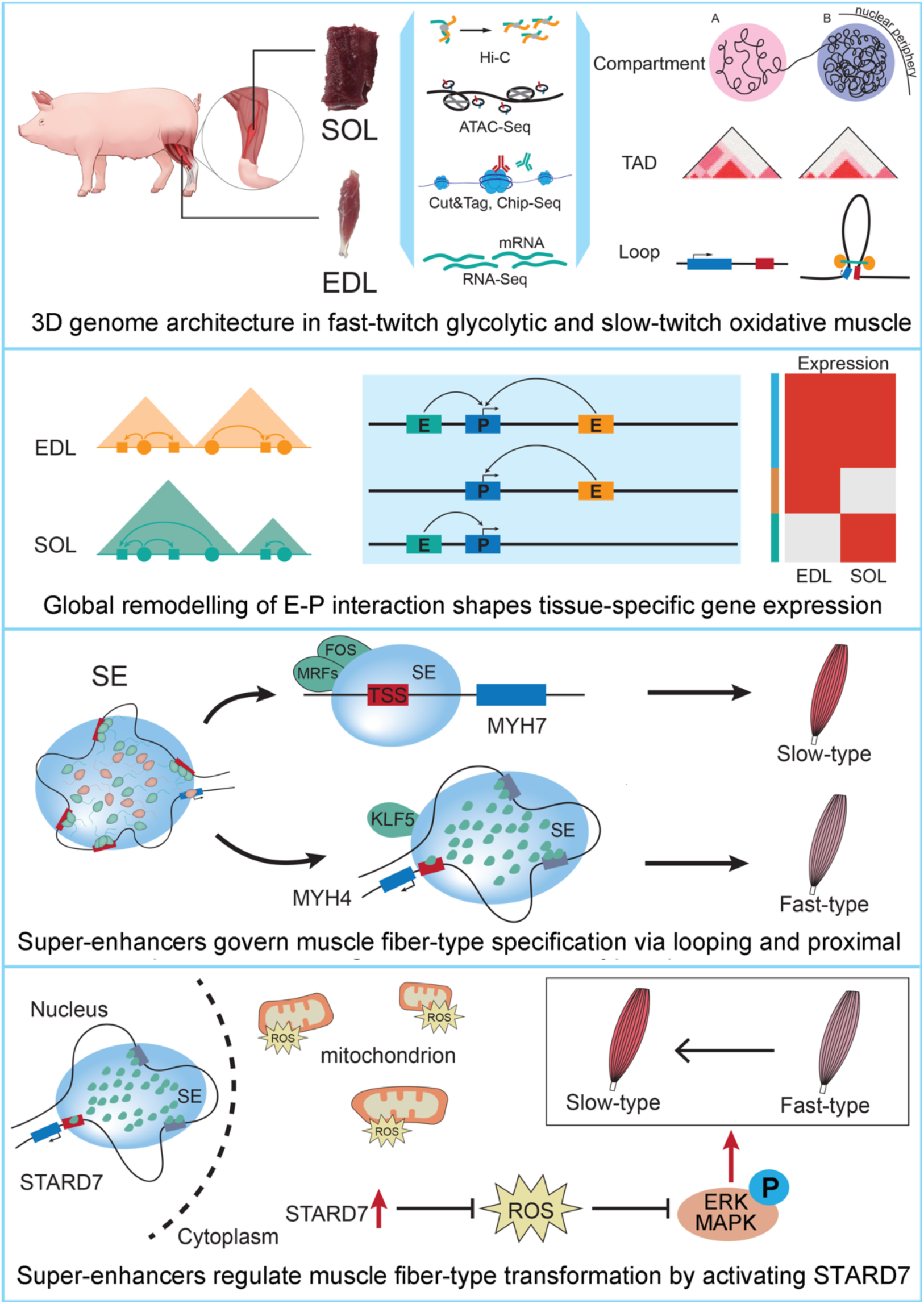

## Introduction

Skeletal muscle comprises 40–50% of total body mass in humans and plays a pivotal role in insulin-stimulated glucose disposal, making it essential for maintaining systemic glucose homeostasis. Dysregulated skeletal muscle metabolism impairs insulin sensitivity and contributes to metabolic disorders such as obesity and Type 2 diabetes (*1, 2*). Muscle fibers exhibit metabolic diversity, with distinct properties shaped by their plasticity and fiber-type composition. Myofibers are classified based on the expression of myosin heavy chain (MyHC) isoforms: slow-twitch fibers predominantly express MYH7 (MyHC I), while fast-twitch fibers express MYH1 (MyHC IIa) and MYH4 (MyHC IIb) (*3*). Slow fibers favor oxidative metabolism and endurance, whereas fast fibers rely on glycolysis and contract more rapidly (*4*).

Although the total number of muscle fibers remains stable in adults (*2*), fiber-type composition can adapt to external stimuli such as aging, exercise, or disease. In obesity and Type 2 diabetes, a shift from oxidative slow fibers to glycolytic fast fibers contributes to impaired insulin sensitivity, reduced metabolic flexibility, and insulin resistance (*5*). In livestock production, the slow-fiber-rich soleus (SOL) muscle offers superior meat quality, characterized by higher redness, intramuscular fat, water-holding capacity, and lower tenderness, compared to the fast-fiber-rich extensor digitorum longus (EDL) muscle (*6, 7*). Understanding the regulatory mechanisms underlying muscle fiber specification and transformation could provide therapeutic insights for human metabolic diseases and strategies to enhance meat quality.

Advances in genome-wide 3D chromatin interaction mapping have revolutionized our understanding of spatial genome organization and its regulatory impact on gene expression. The mammalian genome is hierarchically structured into chromatin loops, topologically associating domains (TADs), and A/B compartments within the nucleus (*8*). TAD boundaries, primarily defined by the CCCTC-binding factor (CTCF) and cohesin complexes (*9*), are largely conserved across cell types and species (*10*). However, chromatin loops and epigenetic landscapes within TADs exhibit cell-type specificity, driving the precise spatiotemporal expression of lineage-specific genes (*11*). Proper 3D chromatin architecture is critical for transcriptional regulation during processes such as myogenesis (*12*), prenatal muscle development (*13*), and muscle stem cell aging (*14*). These findings underscore the importance of 3D genome remodeling in shaping transcriptional programs during muscle fiber differentiation.

*cis*-regulatory elements (CREs), including enhancers and promoters, are defined by specific epigenetic features and regulate gene expression through cell-type-specific chromatin looping. Super-enhancers (SEs), large enhancer clusters, are key regulators of lineage-specific gene expression (*15*). SEs coordinate with chromatin regulators to form extensive intra-TAD loops, enabling precise gene activation (*16*). Previous studies revealed enhancer repertoires in distinct muscles, and highlighting the crucial role of transcription factors (TFs) in fiber-type specification (*17*). Moreover, chromatin looping bound by SEs has been shown to activate fast-type myosin gene expression (*18*). However, the genome-wide dynamics of enhancer-promoter (E-P) interactions between muscles, particularly their role in shaping muscle-specific phenotypes, remain unclear.

The pig (*Sus scrofa*) serves as a valuable biomedical model for studying energy metabolism and muscle diseases due to its physiological, anatomical, and metabolic similarities to humans (*19*). In this study, we provide a comprehensive multi-omics dataset from pig EDL and SOL muscles to clarify the regulatory mechanisms underlying their developmental and morphological differences. We identified numerous active CREs and demonstrated how tissue-specific E-P interactions and TFs orchestrate transcriptional programs in distinct muscles. Furthermore, we identified key SEs that regulate slow- and fast-type myosin gene expression via chromatin interactions. Notably, we validated that SE-driven activation of *STARD7* induces the transformation of glycolytic to oxidative fibers by reducing reactive oxygen species (ROS) and inhibiting ERK MAPK signaling. Together, our findings highlight the critical role of activated CREs and 3D genome organization in muscle fiber specification and transformation.

## Results

### Transcriptome dynamic and genome-wide analysis of CREs in SOL and EDL

In this study, we collected SOL and EDL muscles from three 6-month-old Duroc pigs to compare their muscle characteristics (Fig. 1A). Slow-twitch fibers predominated in the SOL, while fast-twitch fibers were more abundant in the EDL, as evidenced by differential expression of myosin genes (Fig. 1B-C; Fig. S1A-C). RNA-seq analysis of SOL and EDL muscles identified 788 differentially expressed genes (DEGs) including 366 SOL-induced DEGs and 412 SOL-induced DEGs (Fig. S1D; table S1). Gene Ontology (GO) analysis and Gene Set Enrichment Analysis (GSEA) further revealed that glycolysis and muscle contraction pathways were more active in the EDL-induced DEGs, while oxidative phosphorylation and fatty acid β-oxidation pathways were more prominent in the SOL-induced DEGs (Fig. S1E-F; table S1). To analyze the rationality of using bulk tissues to present the status of myonuclei, we compared the epigenetics signal of bulk tissue and purified-myonuclei. We found that genome-wide H3K27ac signal showed a high similarity (R>0.8) between bulk tissue and purified-myonuclei (Fig. S2A-B). The majority of H3K27ac peaks (over 80%) identified in myonuclei are common in corresponding bulk tissue (Fig. S2C). These findings confirmed that SOL and EDL can be respectively classified as typical oxidative and glycolytic muscles, and could greatly reflect the epigenetic statues of purified-myonuclei, making them ideal for studying the regulatory mechanisms underlying muscle fiber-type specification and transformation.

**Fig. 1.**
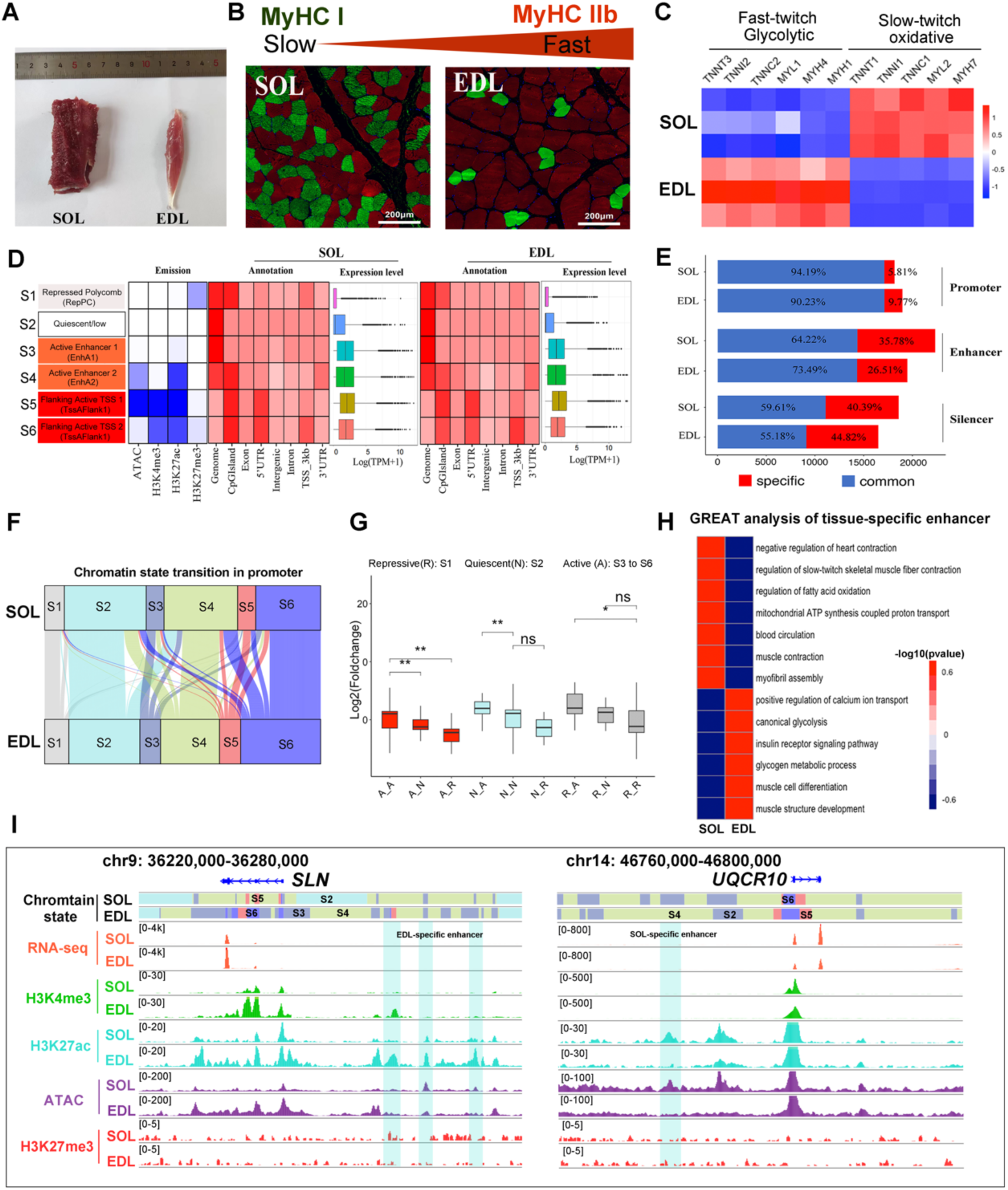
Transcriptome dynamics and genome-wide analysis of CREs in SOL and EDL. (A) Isolated EDL and SOL muscles. (B) Representative immunofluorescent staining of slow-twitch MyHC I (red) and fast-twitch MyHC IIb (green) in SOL and EDL sections. Magnification: ×20, Bar = 200 μm. (C) Heatmap of muscle fiber-type marker gene expression. (D) Classification and annotation of chromatin states, with each row representing a chromatin state and each column denoting histone marks, chromatin accessibility, genomic annotation, and gene expression levels. TssAFlank1: high H3K4me3 and ATAC-seq signal; TssAFlank2: high H3K4me3 but weak ATAC-seq signal. EnhA1: high H3K27ac and ATAC-seq signal; EnhA2: weak H3K27ac and ATAC-seq signal. (E) Tissue specificity of identified CREs. (F) Alluvial diagram depicting changes in promoter chromatin states of DEGs. (G) Expression changes of DEGs with switched chromatin states. (H) Functional enrichment analysis of tissue-specific enhancers using GREAT. (I) ATAC-seq tracks, chromatin states, and histone modifications at *SLN* and *UQCR10* loci and their associated enhancers in SOL and EDL.

To investigate the relationship between chromatin epigenetic states and transcriptional regulation, we conducted ChIP-seq for H3K27me3, CUT&Tag for H3K27ac and H3K4me3, and Assay for Transposase-Accessible Chromatin (ATAC-seq) to assess chromatin accessibility in both SOL and EDL muscles. Principal component analysis (PCA) of each epigenetics mark datasets demonstrated high reproducibility between biological replicates (Fig. S3A; table S2). Profiles of histone modifications and chromatin accessibility revealed that transcriptional activation was associated with increased chromatin accessibility and enrichment of active histone marks (H3K27ac and H3K4me3) (Fig. S3B).

We applied ChromHMM to these epigenetics datasets, and categorized the six distinct chromatin states, each associated with specific gene expression patterns (Fig. 1D). The active state consisted of active transcription start site (TSS) proximal promoter states (TssFlank1 and TssFlank2), enhancer-related states (EnhA1 and EnhA2). The repressed polycomb (ReprPC) state enriched with H3K27me3 are defined as repressive state. The remaining of genome was denoted as quiescent state with low epigenetics mark signals. Based on the genome state classification, we identified 18,610–19,961 candidate promoters, 21,081–24,128 candidate enhancers, and 16,473–18,564 candidate silencers in SOL and EDL muscles (Fig. 1E; table S3). The transition of promoter chromatin states from repressive or quiescent to active was accompanied with the activation of gene transcription, which was associated with the tissue-specific activation of glycolysis-related and contraction-related genes in SOL and EDL, such as *BPGM* and *CACNA2D3* (Fig. 1F-G; Fig. S3C).

We considered the tissue specificity of candidate enhancers, and observed that over 26% enhancers show differential activity between SOL and EDL muscles (Fig. 1E). Comparative analyses across pig, human, and mouse genomes revealed significant differences in enhancer activity (Fig. S3D). Genomic Regions Enrichment of Annotations Tool (GREAT) enrichment analysis indicated that tissue-specific enhancers were located near genes critical for muscle contraction and metabolic processes (Fig. 1H; Fig. S3E). For instance, EDL-specific enhancers were found near the *SLN*, which regulates calcium transport essential for fast-twitch muscle contraction (*20*). Similarly, SOL-specific enhancers were identified near the *UQCR10*, which is involved in the mitochondrial respiratory chain and oxidative phosphorylation (Fig. 1I) (*21*). In summary, our results demonstrated that dynamic epigenetic states of promoters and enhancers may play pivotal roles in regulating transcriptomic changes associated with muscle fiber-type development.

### Characterization of the 3D Chromatin Organization in SOL and EDL

The distinct epigenetic landscapes of SOL and EDL muscles highlight the importance of chromatin remodeling in defining oxidative and glycolytic muscle phenotypes. To investigate the multi-scale reorganization of chromatin structure and its effect on gene expression, we employed Bridge Linker-Hi-C (BL-Hi-C) to generate genome-wide chromatin interaction maps in SOL and EDL muscles. The Hi-C datasets of biological replicates with high correlation were merged, resulting in ~2.31 billion uniquely valid pairwise contacts (Fig. S4A-B; table S2). These datasets achieved an ultra-deep resolution of 5 kilobase pairs (kb), enabling a detailed analysis of 3D genome organization (Fig. S4C-D).

We analyzed genome compartmentalization into A and B compartments at a 50 kb resolution. Regions in the A compartment were characterized by higher GC content, greater gene density, elevated gene expression, and increased levels of active histone modifications and chromatin accessibility compared to regions in the B compartment (Fig. S4E; table S4). Compartment segregation was largely conserved, with only ~4.67% of the genome undergoing compartment switching between SOL and EDL (Fig. 2A). These compartment changes were associated with gene expression shifts: regions transitioning from B to A compartments showed increased expression, while A to B transitions corresponded to decreased expression (Fig. 2B). Notably, the transcription factor SIM1, crucial for energy balance and obesity resistance (*22*), showed increased expression in EDL in conjunction with a B to A compartment switch (Fig. 2C).

**Fig. 2.**
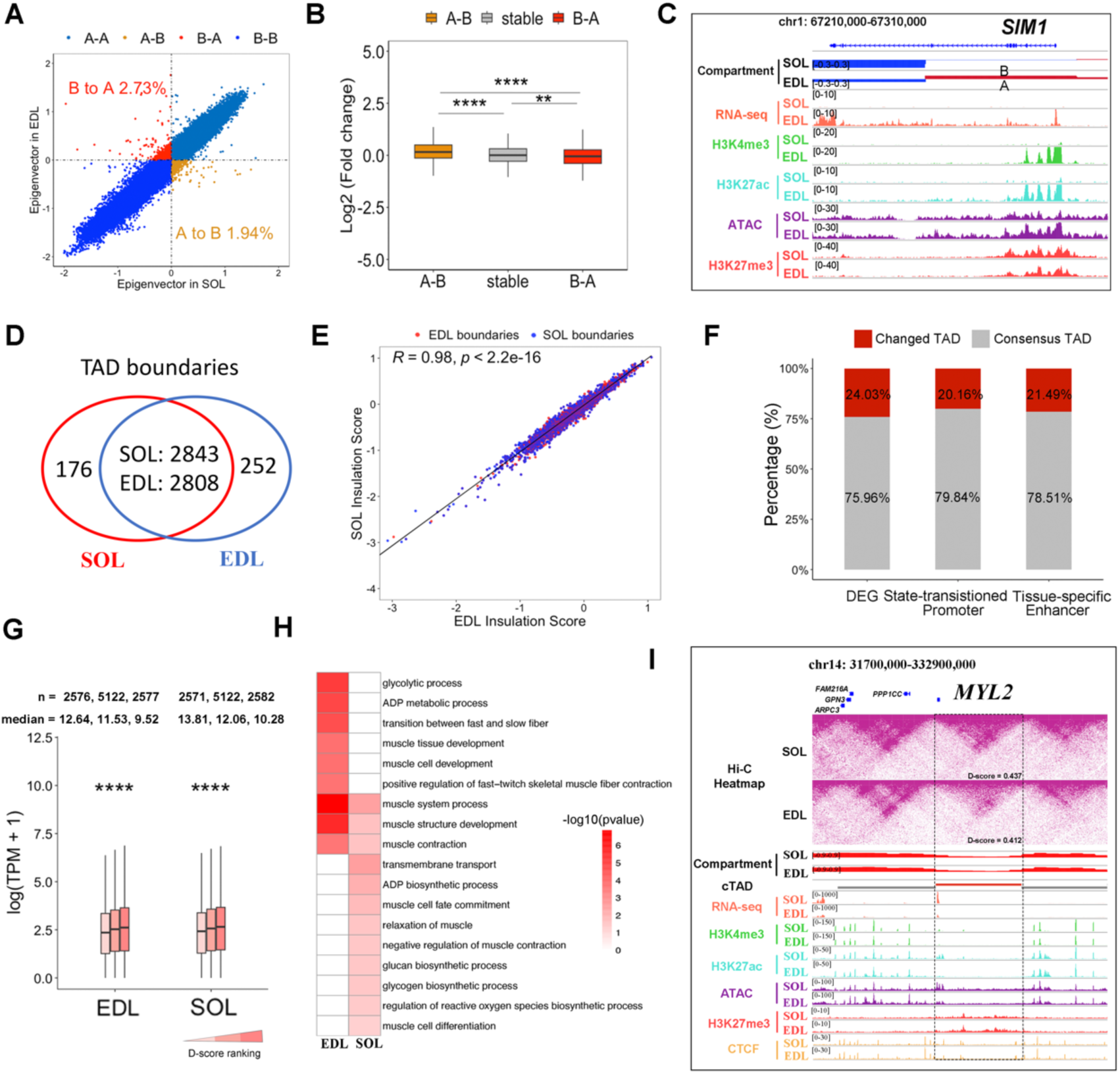
Characterization of the 3D Chromatin Organization in SOL and EDL. (A) Compartment switching between SOL and EDL. (B) The expression changes of genes located within regions that underwent the compartment switch (A-to-B: *n* = 576; B-to-A: n=536; stable: n=11581). The horizontal line denotes the median, the box encompasses the interquartile range, and the whiskers extend to the 5^th^ and 95^th^ percentiles. (C) *SIM1* gene locus underwent a B-to-A switch from SOL to EDL, with higher expression in EDL than SOL. Tracks for RNA-seq and ATAC-seq signal, and histone modification are shown. (D) Overlap of TAD boundaries between SOL and EDL. (E) Correlation of insulation scores for SOL and EDL; Pearson correlation coefficient is indicated (*R* = 0.98). (F) The distribution of DEGs and CREs in changed TADs and cTADs. (G) Gene expression levels in cTADs (*n* = 2376) with relatively low, medium, or high D-scores. (H) The enriched GO terms for genes within the cTADs with higher D-scores in EDL (EDL) and cTADs with higher D-scores in SOL (SOL). (I) Representative cTAD with different D-scores in two tissues. Top: Hi-C contact heatmaps of the genomic region containing *MYH2* gene locus. D-scores of cTAD were marked. Middle: TAD boundaries and genome browser tracks of PC1 values. Bottom: tracks for RNA-seq and ATAC-seq signal, CTCF, and histone modification. \*\**P* < 0.01, \*\*\*\**P* < 0.0001, two-sided Wilcoxon test.

We identified 2,999 TADs (3,018 TAD boundaries) in SOL and 3,041 TADs (3,060 TAD boundaries) in EDL, with only 5.82–8.16% of TAD boundaries being tissue-specific (Fig. 2D-E; Fig. S4F; table S5). The accuracy of TAD boundaries was validated using directionality index (DI) and insulation score (IS) analyses (Fig. S4G). As expected, CTCF was enriched at TAD boundaries and contributed to TAD insulation strength (Fig. S4H-I). Genome-wide comparison of insulation scores revealed high conservation of TAD structures between SOL and EDL, with a strong correlation (*R* = 0.98) and similar insulation profiles (Fig. 2E).

To assess intra-TAD interactions, we identified consensus TADs (cTADs) whose boundaries are stable in SOL and EDL and accounted for 73.50% of the genome. We found that DEGs and CREs with dynamic activity were primarily located within cTADs (Fig. 2F). To quantify cTAD’s tendency to self-interact, we calculated domain score (D-score) defined by the ratio of intra-TAD contacts to all cis contacts within a 2 Mb window (Fig. S4J; table S5). D-scores positively correlated with the expression levels of genes within cTADs (Fig. 2G). GO enrichment analysis of genes within cTADs revealed that dynamic intra-TAD connectivity influenced essential genes for muscle contraction and metabolism (Fig. 2H). For instance, myosin light chain-2 (*MYL2*), a gene critical for myosin ATPase activity and predominantly expressed in slow-twitch fibers, exhibited both higher expression and D-scores in SOL compared to EDL (Fig. 2I).

In conclusion, while compartmentalization and TAD structures were largely conserved between SOL and EDL muscles, intra-TAD interactions showed significant changes and correlated with dynamic transcriptional activity.

### Global remodeling of E-P interactions underpins functional divergence in SOL and EDL

To uncover the role of distal chromatin interactions among CREs in gene regulation, we conducted loop calling to identify chromatin loops in SOL and EDL muscles. This analysis revealed 16,086 chromatin loops in SOL and 12,923 in EDL (Fig. 3A; table S6), with most loops occurring within TAD boundaries (Fig. 3B). Chromatin loops were categorized based on CREs at their anchors, with E-P interactions constituting the largest proportion in both muscles (Fig. 3C). Aggregate peak analysis (APA) of different loop subtypes demonstrated that loop interaction strength varied by subtype and was positively correlated with CTCF enrichment, underscoring CTCF’s critical role in loop formation (Fig. S5A-C). Motif analysis further identified significant enrichment of MEF2, KLF, and AP-1 TF families in different loop subtypes (Fig. S5D-E).

**Fig. 3.**
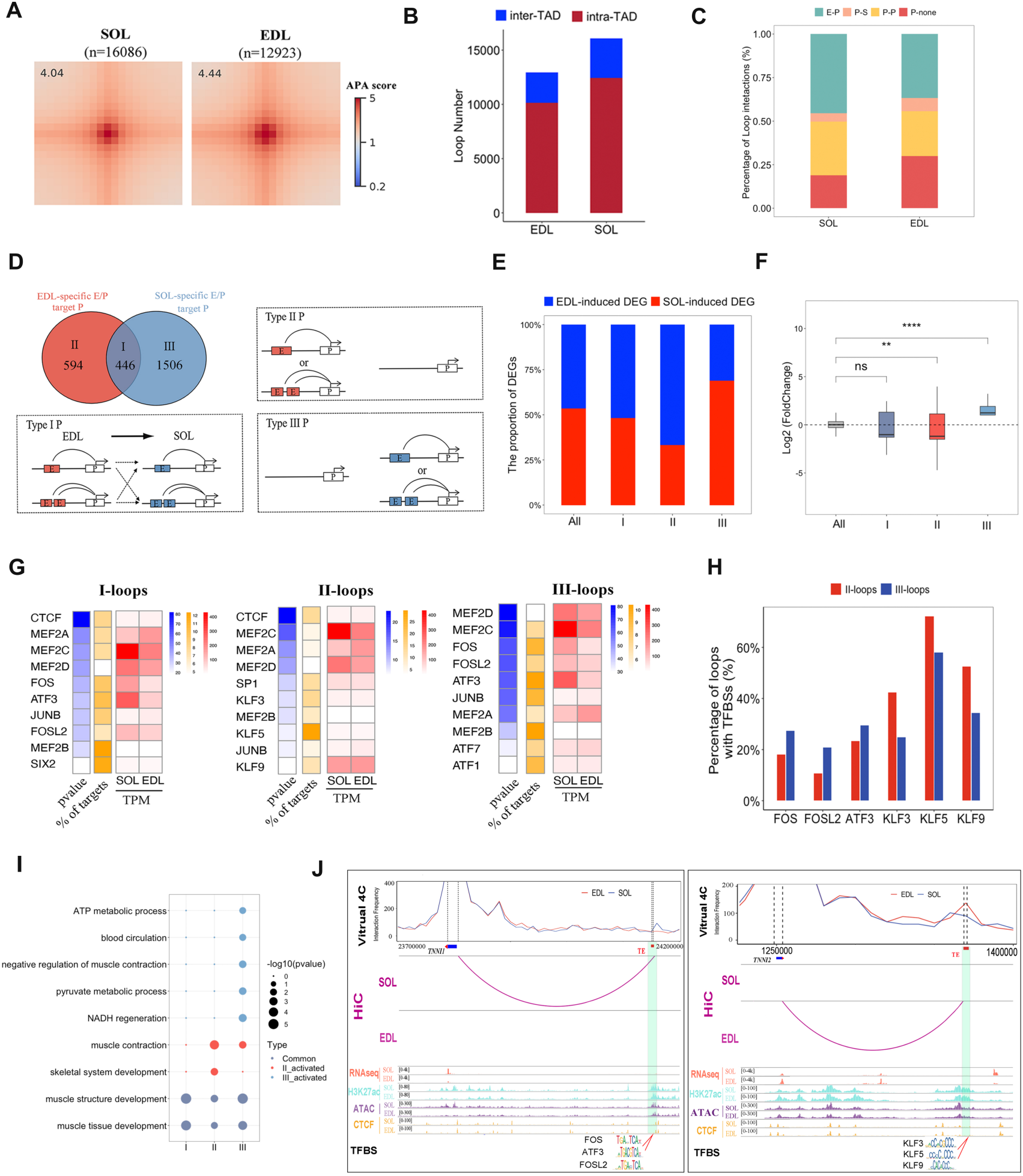
Global remodeling of E-P interactions underpins functional divergence in SOL and EDL. (A) APA analysis of chromatin loops in SOL (*n* = 16,086) and EDL (*n* = 12,923). Bin size: 10 kb. (B) Proportion of inter-TAD and intra-TAD loops. (C) Distribution of loop interaction types: P-S (promoter-silencer), P-P (promoter-promoter), P-E (promoter-enhancer), and P-none (promoter with no other CRE). (D) Overlap of genes with distinct E-P interactions (top-left). Schematic representation of E-P rewiring for Groups I-III. (E) The proportion of DEGs in Groups I-III. (F) Transcriptional changes for genes in Groups I-III. Log2 (Foldchange) represents as the change of gene expression in SOL compared to EDL. (G) Top 10 enriched TFs in Groups I-III. (H) Percentage of loops with TFBS for selected TFs. (I) The enriched GO terms for genes in Group I-III. II_activated: GO term that more activated in GroupII; III_activated: GO term that more activated in Group III; Common: GO term common in three Groups. (J) Chromatin interactions around *TNNI1* and *TNNI2*. Top: Virtual 4C profile of loop contact differences. Tracks for chromatin loops, RNA-seq, ATAC-seq, CTCF, histone modifications, and TFBS are shown below. Enhancers are marked in green.

To explore the functionality of loops, we compared the expression of genes with different loop subtypes, revealing that the genes with E-P interaction exhibited higher expression than those regulated by other CREs (Fig. S5F). Approximately 86% of enhancers interacted with promoters other than the closest one (Fig. S5G), and multiple enhancers exerted additive effects on gene transcription (Fig. S5H). These findings highlight E-P interactions pivotal role in transcriptional regulation. Thus, we further classified enhancer target genes into three groups (I-III) based on distinct E-P rewiring patterns (Fig. 3D). Genes with SOL-specific interactions (Group III) were predominantly upregulated in SOL (69%, 69 of 100 DEGs), while those with EDL-specific interactions (Group II) were largely upregulated in EDL (66.7%, 24 of 36 DEGs) (Fig. 3E-F). These results suggested that the formation of E-P interactions was associated with the tissue-specific gene activation in SOL and EDL. Motif analysis of these looped enhancers revealed that AP-1 TFs (e.g., *FOS*, *ATF3*, *FOSL2*) were enriched in SOL-specific (Group III) loops, while KLF TFs (e.g., *KLF3*, *KLF9*, *KLF5*) were enriched in EDL-specific (Group II) loops (Fig. 3G). ATAC-seq footprint analysis supported these findings by identifying TFBS corresponding to these TFs (Fig. 3H). We also collected public TFs ChIP-seq datasets of C2C12 myotubes, and observed the enrichment of these TFs in the conserved region of enhancer (Fig. S5I). These findings supported the binding of TFs in identified enhancers. To determine the functional processes of E-P interaction, we applied GO enrichment analysis to Group I-III genes, and found that Group I genes were involved in muscle structure development, while Group II and III genes were more enriched in muscle-contraction- and oxidative-metabolism-related processes, respectively (Fig. 3I). An example of tissue-specific E-P interactions is seen in the troponin complex, essential for the regulation of muscle contraction (*23*). In this study, *TNNI1* (troponin I1, slow-type) and *TNNI2* (troponin I2, fast-type) were associated with SOL-specific and EDL-specific E-P interactions, respectively, aligning with their expression patterns (Fig. 3J).

In summary, our findings suggest that AP-1 and KLF families may contribute to the formation of SOL- and EDL-specific E-P interactions, respectively. This global remodeling of E-P interactions may drive tissue-specific transcriptional activation, shaping the distinct contractility of oxidative and glycolytic muscle types.

### Super-enhancers control muscle fiber-type specification via proximity and chromatin looping

SEs are critical in driving lineage- and development-specific gene expression. To explore this, we utilized H3K27ac CUT&Tag datasets to identify SEs. A total of 554 and 592 SEs were identified in the SOL and EDL muscles, respectively (Fig. 4A, table S4). The majority of SEs were located in compartment A and within cTADs (Fig. S6A). We compared genes with promoter overlapped with SEs (SE-proximal genes) in SOL and EDL muscles, and identified 177 EDL-specific SE-proximal genes and 140 SOL-specific SE-proximal genes. To elucidate the function of SEs, we categorized SE-proximal genes based on whether their promoters overlapped with SEs in SOL and EDL muscles, revealing that SEs in gene promoters might activate tissue-specific gene transcription (Fig. S6B). GO analysis of SE-proximal genes demonstrated that SOL-specific and EDL-specific SE-proximal genes were respectively more activated in muscle contraction and oxidative metabolism (Fig. 4B), underscoring SEs’ roles in functional divergence between SOL and EDL muscles. Among these, a SOL-specific SE located in the *MYH7* promoter (designated SE-MYH7) exhibited strong H3K27ac signal and chromatin accessibility, suggesting its potential role as a key regulatory element for *MYH7* activation in slow-twitch SOL muscle (Fig. 4C). TF footprint analysis of differential ATAC peaks at SE-MYH7 revealed the potential binding of TFs, included *MyoG* (*24*), *Hoxa13* (*25*), and *Vdr* (*26*) those have been reported to bind on the *MYH7* promoter or activate *MYH7* expression, as well as unreported TFs including FOS (AP-1 family). FOS was highly expressed in SOL, and predicted to bound in most SOL-specific enhancers, suggesting its crucial role in activating SE-MYH7.

**Fig. 4.**
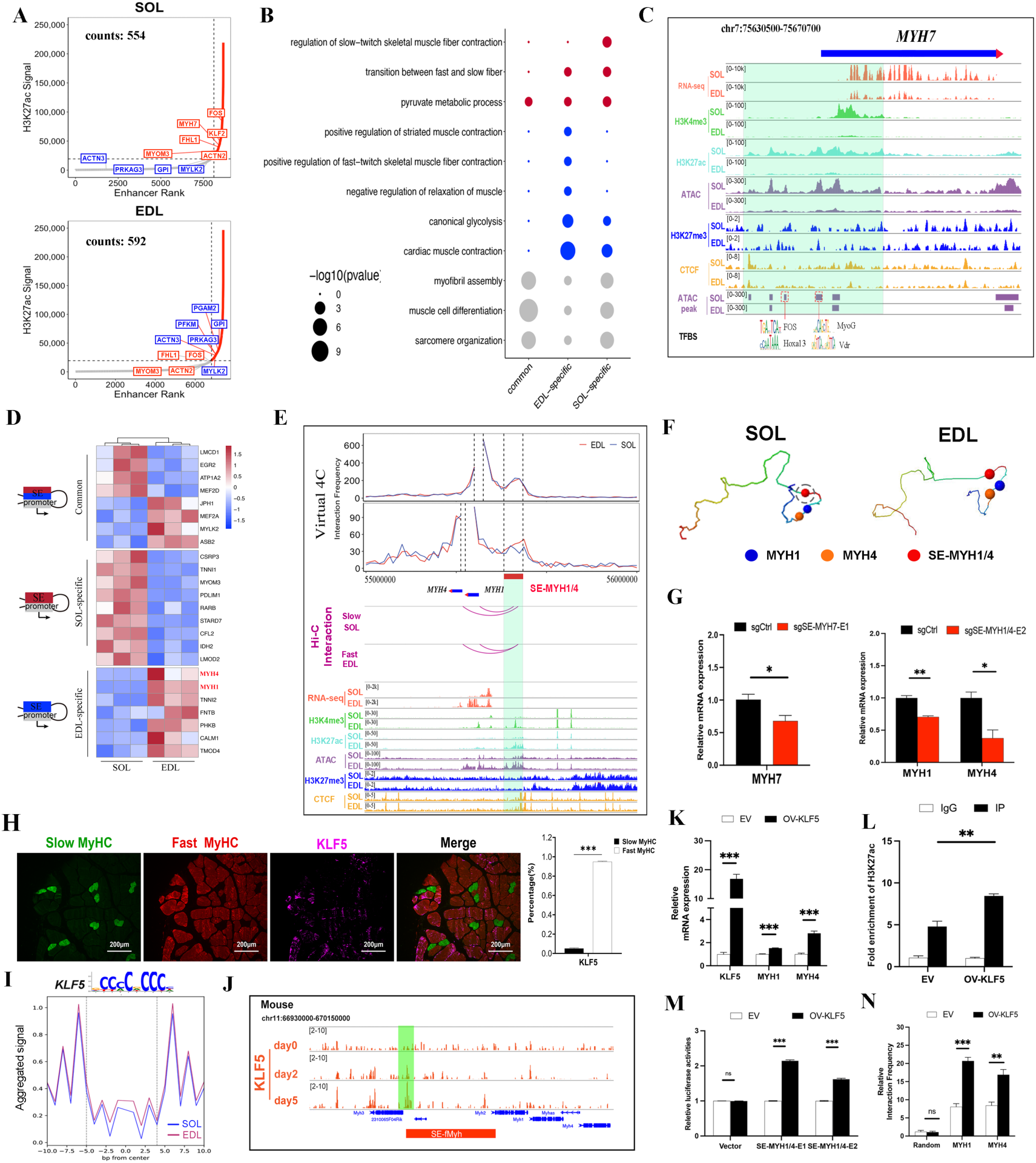
Super-enhancers control muscle fiber-type specification via proximity and chromatin looping. (A) Hockey stick plots based on input-normalized H3K27ac signals in SOL and EDL, highlighting SOL-specific genes (red) and EDL-specific genes (blue). (B) GO term enrichment for SE-proximal genes. (C) Tracks of RNA-seq, ATAC-seq, CTCF, histone modifications, and TFBS at the *MYH7* locus with SE regions indicated. (D) Expression heatmap of SE-anchored genes. (E) Epigenetic states and chromatin interactions at *MYH1*/*MYH4*. Virtual 4C plots show chromatin contact differences; tracks for RNA-seq, ATAC-seq, CTCF, and histone modifications are included. SEs are marked in green. (F) 3D chromatin conformation models around the gene locus (bin size: 10 kb). (G) *MYH7* and *MYH4* expression changes after SE deletion via CRISPR-Cas9 in PSCs. (H) Immunofluorescence staining of KLF5, MYH4 (Fast-MyHC) and MYH7 (Slow-MyhC) in the EDL muscle. (Green: MYH7; Red: MYH4; Purple: KLF5). Percentage: The proportion of KLF5 signal in the slow-MyHC or fast-MyHC. (I) Aggregate TF footprint plots for KLF5 in SOL and EDL. (J) Track of KLF5 ChIP-seq in C2C12 myotubes differentiated for 0 day, 2 days, and 5 days. KLF5 peaks was filled in green. (K) qPCR showing the expression change after KLF5 overexpression. (L) ChIP-qPCR showing the enrichment of H3K27ac in SE-MYH1/4 after KLF5 overexpression in differentiated PSCs. (M) Dual-reporter assay detecting the enhancer activity of SE-MYH1/4 after KLF5 overexpression in differentiated PSCs. (N) 3C-qPCR showing the differences of interaction frequencies between SE-MYH1/4 and gene promoters in SOL and EDL. \**P* < 0.05, \*\**P* < 0.01, \*\*\**P* < 0.001, two-sided unpaired *t-*test.

To clarify the functionality of SE depending on distal looping, we annotated SEs to nearest loop anchors. In the SOL and EDL muscles, we identified 1,352 and 817 SE-anchored loops, respectively, corresponding to 365 and 673 SE-promoter interactions (Fig. S6C). Among 46 SE-anchored DEGs, the fast-type myosin *MYH1* and *MYH4* were found to interact with an EDL-specific SE (designated SE-MYH1/4) via chromatin looping (Fig. 4D). Virtual 4C profiling and 3D chromatin modeling further supported these interactions (Fig. 4E-F).

To assess conservation across species, we analyzed H3K27ac ChIP-seq datasets from mouse EDL and SOL (*17*). A conserved SE near the *Myh3* gene (SE-fMyh) was identified, which interacts with and activates fast-type myosin genes, consistent with previous findings (*18*). Approximately 50% of H3K27ac peaks within pig SE-MYH1/4 were sequence-conserved with SE-fMyh, with over 60% sequence identity (Fig. S6D). However, the clustered enhancers in the *myh7* promoter of mice were not classified as SEs by the ROSE algorithm (*27*), despite exhibiting strong H3K27ac enrichment and chromatin accessibility (Fig. S6D). To experimentally validate the functionality of theses SEs, we transfected the pGL3-enhancers vector into differentiated porcine satellite cells (PSCs), and confirmed their enhancer activity by performing Dual-luciferase reporter assays (Fig. S6E). We also deleted conserved H3K27ac peaks within SE-MYH1/4 and SE-MYH7 in PSCs using CRISPR-Cas9, resulting in significant reductions in *MYH1*/*MYH4* and *MYH7* expression, respectively (Fig. 4G; Fig. S6F-G).

To explore the coordinated regulation of TFs with SE-MYH1/4, we ranked expressed TFs based on proportion of SEs with corresponding TFBS, revealing that KLF family as likely key regulators of SEs in EDL (Fig. S6H). The gene expression of most KLFs was expectedly higher expressed in EDL than SOL (Fig. S6I). Thus, we focused on KLF family binding at SE-MYH1/4, which is specifically activated in EDL. We validated that KLF5 is the most differentially expressed KLF transcription factors between SOL and EDL muscles, and higher expressed in fast-type fibers than in slow-type fibers (Fig. 4H). ATAC-seq footprint analysis indicated that KLF5 displayed distinct binding patterns, with higher binding scores in EDL compared to SOL (Fig. 4I; Fig. S6J-K; table S7). Furthermore, we observed a significant enrichment of KLF5 in the SE-fMyh in C2C12 myotubes differentiated for 0 day, 2 days, and 5 days (Fig. 4J). These results suggested that KLF5 might be the key TFs to regulate the activity of SE-MYH1/4 in EDL. To reveal the regulatory mechanism of KLF5 and SE-MYH1/4, we validated the binding of KLF5 and SE-MYH1/4 in EDL by ChIP-qPCR (Fig. S6L). Additionally, we demonstrated that KLF5 induces the expression of *MYH1* and *MYH4* genes by increasing enrichment of H3K27ac and enhancer activity of SE-MYH1/4, and enhances the interaction frequency of SE-MYH1/4 with gene promoter (Fig. 4K-N). These results demonstrated that KLF5 can regulate the enhancer activity and interaction structure of SE-MYH1/4 to induce the formation of fast-type fiber.

### The function of super-enhancers in muscle fiber-type transformation

In this study, we observed that 742 SE-anchored genes are significantly enriched in processes related to muscle fiber-type transformation, including oxidative phosphorylation and calcium ion transport (Fig. S7A). To identify candidate SE-anchored genes essential for this transformation, we calculated the expression correlation of 46 SE-anchored DEGs with *MYH4* and *MYH7* by analyzing transcriptomic datasets of porcine skeletal muscle across eight developmental stages (*28*). From this analysis, 23 SE-regulated DEGs were identified as candidate genes displaying opposite correlations in expression with *MYH4* and *MYH7* (|*R*| > 0.5) (Fig. 5A).

**Fig. 5.**
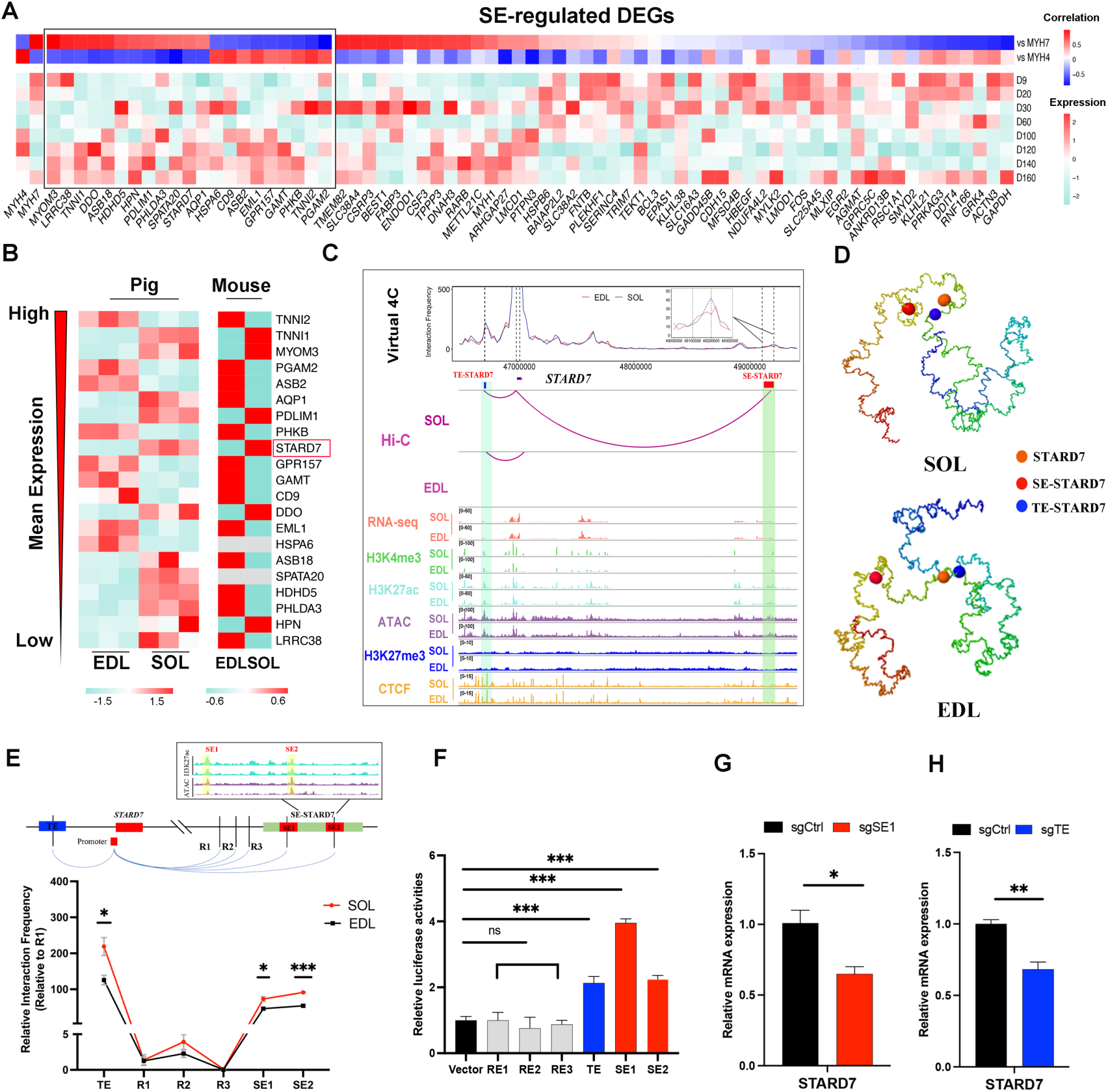
The function of super-enhancers in muscle fiber-type transformation. (A) Top: Pearson correlation coefficients between SE-regulated DEGs and *MYH7* or *MYH4*. Bottom: Expression heatmap of SE-regulated DEGs in postnatal pig skeletal muscle at eight developmental stages. (B) Heatmap of 22 SE-regulated DEGs conserved between pig and mouse SOL and EDL. (C) Virtual 4C plot of chromatin interactions around the *STARD7* locus. Enhancers are marked in green. (D) 3D chromatin models of the *STARD7* locus based on Hi-C data. (E) 3C-qPCR analysis of *STARD7* promoter interactions in SOL and EDL. Interaction frequency is calculated relative to the R2 site. (F) Dual-luciferase reporter assay validating enhancer activity of TE-STARD7 and SE-STARD7. (G) Expression changes in *STARD7* following TE-STARD7 and SE-STARD7 deletion using CRISPR-Cas9. \**P* < 0.05, \*\**P* < 0.01, \*\*\**P* < 0.001, two-sided unpaired *t-*test.

We further examined DEGs with conserved expression patterns in pig and mouse EDL and SOL by accessing MuscleDB (Fig. 5B) (*29*). Among of these candidate genes, *STARD7* is reported as a key gene for myoblast differentiation in mice and humans (*30*). Consistent with its higher expression in SOL compared to EDL (Fig. S7B), *STARD7* expression positively correlated with *MYH7* (*R* > 0.57) and negatively with *MYH4* (*R* < −0.61) during muscle development, suggesting its critical role in muscle fiber-type transformation. Validation through Virtual 4C profiling, 3D chromatin modeling, and 3C-qPCR confirmed the interaction of a typical enhancer (TE-STARD7) and a super-enhancer (SE-STARD7) with the *STARD7* promoter (Fig. 5C-E). Dual-luciferase reporter assays further validated the enhancer activity of both TE-STARD7 and SE-STARD7 in differentiated PSCs (Fig. 5F). CRISPR-Cas9 deletion experiments demonstrated reduced *STARD7* expression upon enhancer deletion (Fig. 5G; Fig. S7C). These findings suggest that SEs activate *STARD7* transcription to drive muscle fiber-type transformation.

### *STARD7* induces the transformation of glycolytic fiber to oxidative fiber

To investigate the role of *STARD7* in muscle fiber transformation, we conducted gain- and loss-of-function experiments both *in vitro* and *in vivo*. In PSCs, qPCR, cell immunofluorescence staining, and WB results confirmed that *STARD7* promotes the transformation of glycolytic myotubes to oxidative myotubes in differentiated PSCs. (Fig. 6A-F). We measured extracellular acidification rate (ECAR) to detect glycolysis activity using seahorse assays. Our results validated that ECAR decreases after STARD7-overexpression, while ECAR increased after STARD7-knockdown in living differentiated PSCs (Fig. 5G-H). Additionally, we validated that STARD7 promotes the expression of marker genes of oxidative metabolism and slow-type troponin, while decreases the expression of marker genes of glycolysis metabolism and fast-type troponin (Fig. 5I-J). These results demonstrated that *STARD7* could induce the glycolytic to oxidative metabolism. For *in vivo* validation, we respectively injected LV3-shSTARD7 or LV3-shNC particles into the right and left hindlimbs for 2-months, respectively. We collected Gastrocnemius (GAS) muscles in 8 time points to monitor the expression change of slow-type myosin (MYH7) and fast-type myosin (MYH4) (Fig. 6K). Our results validated that, after 5-week injection, S*TARD7* knockdown reduces the mRNA and protein levels of *MYH7* while increases *MYH4* levels by performing RT-qPCR, WB and tissue immunofluorescence staining (Fig. 6M-O; Fig. S8A). Interestingly, our results indicated that overexpression of pig *STARD7* gene can also be able to induce the fast-type fibers to slow-type fibers in mouse (Fig. S9A-C). In 8-week injection, we validated that *STARD7* knockdown results in increased muscle weight and altered meat color characteristics, including higher L* (lightness) and a* (redness) values (Fig. 6P-Q). Collectively, these results confirmed that *STARD7* can induce glycolytic-to-oxidative fiber transformation, decrease muscle weight, and enhance muscle redness and lightness.

**Fig. 6.**
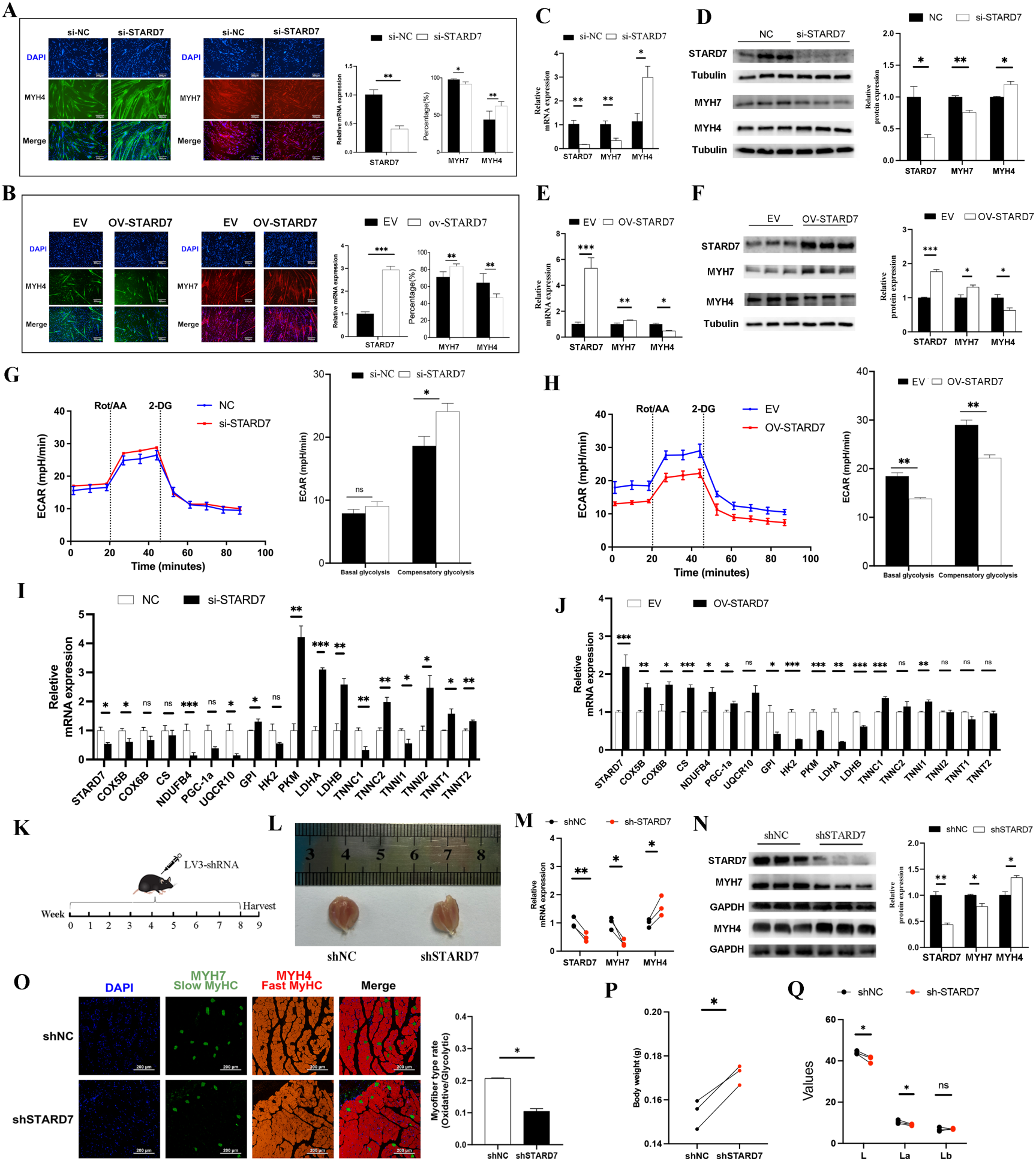
STARD7 induces the transformation of glycolytic fiber to oxidative fiber. (A-B) Immunofluorescence staining of fast-type *MYH4* and slow-type *MYH7* following *STARD7* knockdown (A) and overexpression (B). left panel: representative images of immunofluorescence staining. Middle: expression changes in *STARD7*. Right panel: Statistical analysis of MYH7^+^ and MYH4^+^ fibers. Magnification, × 10, Bar = 200 µm. (C-D) qPCR and WB depicting the gene expression change between *STARD7* knockdown and control group. (E-F) qPCR and WB showing the gene expression change between *STARD7* overexpression and expression vector (EV) group. (G-H) The change of ECAR after STARD7-knockdown (G) and overexpression (H). The average of three independent experiments with four technical replicates for each assay is presented as the mean ± SEM (n = 4 per group). (I-J) The expression changes of maker genes (oxidative: *COX5B*, *COX6B*, *CS*, *NDUFB4*, *PGC-1α*, *UQCR10*; glycolytic: *GPI*, *HK2*, *PKM*, *LDHA*, *LDHB*; slow-type troponin: *TNNC1*, *TNNI1*, and *TNNT1*; fast-type troponin: *TNNC2*, *TNNI2*, and *TNNT2*) after *STARD7* knockdown (I) and overexpression (J). (K) Schematic shSTARD7 or shNC particle injection into GAS muscles and harvesting time points for subsequent analysis. (L) Representative images of GAS muscles from the left (shNC) or right (shSTARD7) muscles. (M-N) qPCR and WB depicting gene expression change between LV3-shSTARD7 and LV3-shNC particles. (O) Immunofluorescence staining of fast-type myosin (MYH4) and slow-type myosin (MYH7) after *STARD7*-knockdown. Left panel: Representative images of immunofluorescence staining. Right panel: statistical analysis of the proportion changes of MYH4^+^ fibers (green) and MYH7^+^ fibers (red). Magnification, × 20, Bar = 50 µm. (P-Q) The comparison of muscle weight and meat color. L: lightness value; a*: redness value; b*: yellowness value. *ns*, no significance, \**P* < 0.05, \*\**P*<0.01, *** *P* < 0.001, two-sided unpaired *t-*test.

### *STARD7* reduces intracellular ROS levels and ERK MAPK activity

STARD7 is a lipid transport protein that delivers phosphatidylcholine to mitochondria (*31*). We validated that the expression of KLFs was not significantly changed after STARD7 overexpression (Fig. S10A). The expression of STARD7 was also not regulated by KLF5 overexpression (Fig. S10B). These results indicated that STARD7 is not involved in the KLFs-mediated regulatory networks. To investigate the regulatory role of *STARD7* in muscle fiber-type transformation, we conducted RNA-seq analysis on *STARD7*-overexpressing (OV-*STARD7*) cells and control cells (Fig. 7A; table S8). The efficiency of *STARD7* overexpression and its ability to facilitate glycolytic-to-oxidative fiber transformation were confirmed (Fig. 7B; Fig. S10C). PCA analysis demonstrated high reproducibility across three biological replicates (Fig. S10D). We identified 215 DEGs after *STARD7* overexpression that enriched in processes related to muscle fiber-type transformation, such as muscle contraction (Fig. 7C; table S8). GSEA results revealed that pathways related to ROS metabolism and ERK-MAPK signaling were downregulated in OV-STARD7 cells (Fig. 7D).

**Fig. 7.**
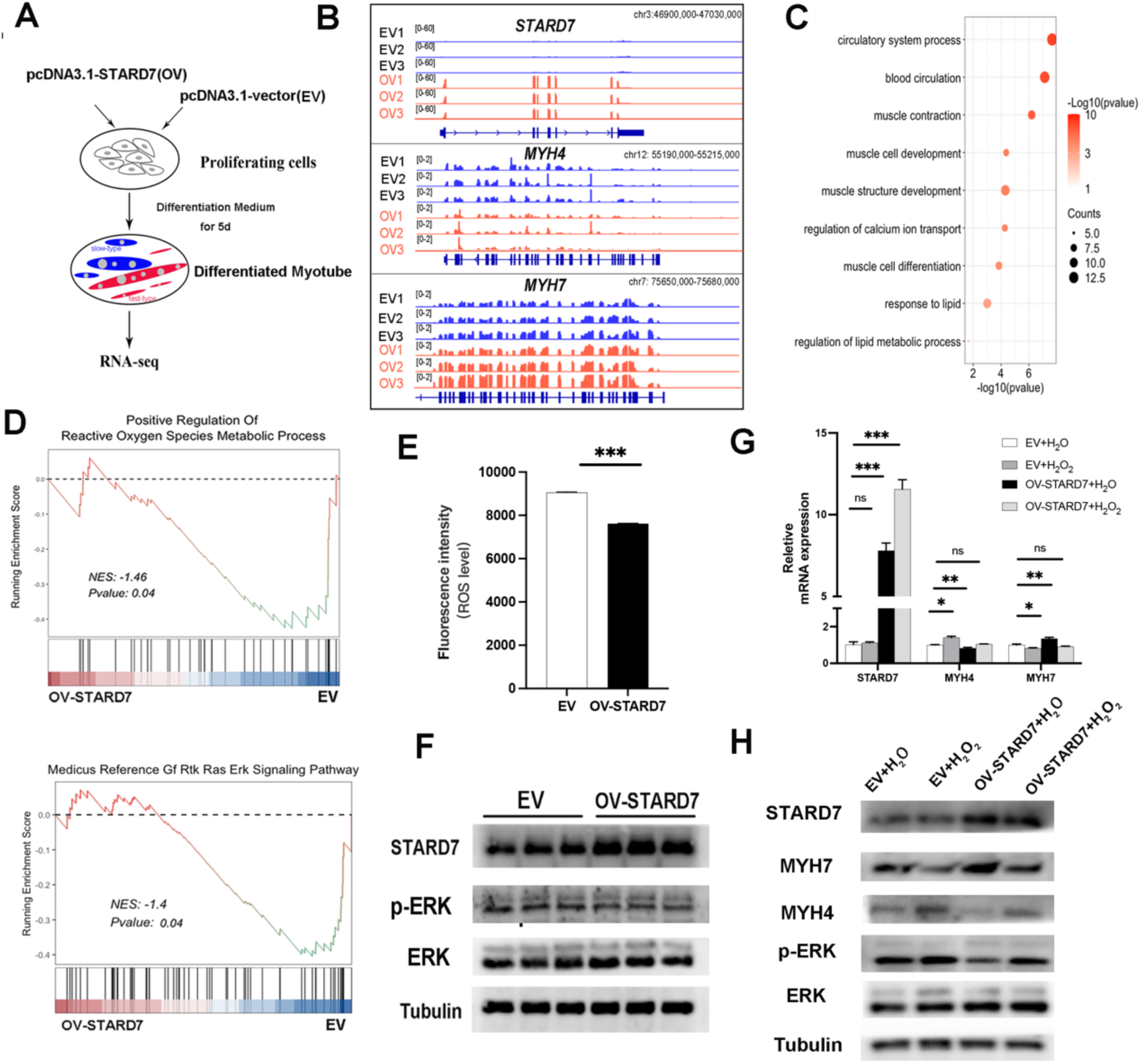
STARD7 reduces intracellular ROS levels and ERK MAPK activity. (A) Schematic of cell culturing and treatment for RNA-seq. (B) RNA-seq tracks for *STARD7*, *MYH4*, and *MYH7* loci in OV-*STARD7* and EV cells. (C) GO term enrichment analysis for DEGs in OV-*STARD7* and control PSCs. (D) GSEA of expressed genes between OV-*STARD7* and EV cells. (E) Measurement of intracellular ROS levels. Excitation wavelength: 488nm, emission wavelength: 525nm. (F) Levels of phosphorylated ERK (p-ERK) and total ERK in OV-*STARD7* and control cells. (G-H) qPCR and WB results showing the impact of exogenous H₂O₂ on STARD7-mediated muscle fiber-type transformation.

Previous studies have shown that *STARD7* reduces ROS levels in HepG2 cells (*32*). Decreased ROS levels suppress MAPK signaling activity in various cell types (*33, 34*), and MAPK activation promotes slow-to-fast fiber transformation in mouse C2C12 myoblasts (*35*). In this study, we validated the preferential accumulation of p-ERK-MAPK in EDL compared to SOL (Fig. S10E). We also evidenced that ROS level is higher in the EDL than SOL by using EMSA assay (Fig. S10F). These results suggested that the reduction of ROS level and p-ERK activity might promote the formation of slow-fiber. To identify the association of ROS level and ERK-MAPK signaling with *STARD7*, we validated a significant reduction in intracellular ROS levels and ERK-MAPK activity in OV-STARD7 cells compared to controls (Fig. 7F). Furthermore, treatment with H₂O₂, which increased ROS levels, disrupted STARD7’s ability to mediate muscle fiber-type transformation, confirming the critical role of ROS in this process (Fig. 7G-H).

## Discussion

Although transcriptional mechanisms underlying muscle fiber-type specification and transformation have been extensively studied, the role of 3D genome architecture reorganization and chromatin state alterations in these processes remains insufficiently understood. In this study, we integrated multi-omics data to uncover how multiscale 3D genome architecture remodeling contributes to transcriptional reprogramming in distinct muscle types. On a large scale, we observed moderate compartment switches and subtle transcriptional changes. While most TAD boundaries remained stable, a subset exhibited dynamic interaction frequencies between muscle types, with reduced intra-TAD interactions correlating with transcriptional inactivation. This highlights the need for further exploration of chromatin loop rewiring within TADs.

By mapping chromatin loops, we precisely linked enhancers to their target genes. We found that Enhancer-promoter (E-P) interactions play a pivotal role in activating genes associated with muscle contraction and oxidative metabolism. For instance, the troponin (Tn) complex, comprising *TNNI1*, *TNNC1*, *TNNI2*, and *TNNT1*, controls Ca²⁺-dependent muscle contraction and adapts to exercise or disuse (*36*). We identified spatiotemporal remodeling of E-P interactions governing Tn complex transcription, underscoring the role of enhancers and chromatin looping in establishing distinct contractile properties across muscles. While TF cooperativity at enhancers is recognized as critical for developmental gene activation, its role in muscle-specific transcription remains poorly understood. Previous studies revealed that AP-1 TFs partner with PGC-1α to regulate hypoxic gene programs in muscle cells (*37*). PGC-1α is a key regulator of metabolism, and can increase oxidative capacity and trigger a shift of fast-type to slow-type fibers (*38*). KLF15 cooperates with transcriptionally factor MRTFB to promote contractile gene expression in vascular smooth muscle cells (*39*). Our findings indicate that the AP-1 and KLF families may shape SOL- and EDL-specific E-P interactions, respectively. Collectively, our study provides a valuable resource for elucidating the relationship between transcriptional reprogramming and 3D genome remodeling, particularly the role of TFs and E-P interactions in tissue-specific gene activation essential for muscle contraction and glucose metabolism.

A key finding of this study is the elucidation of regulatory mechanisms specifying muscle fiber types at the 3D genome level. In adult muscles, the contraction and metabolic properties of specialized myofibers are governed by the expression of slow-type myosin genes (*MYH7*) and fast-type myosin genes (*MYH1, MYH2, MYH4*) (*40*). In mice, fast-type *Myh* genes (*fMyh*) are regulated by SE-*fMyh* through competitive chromatin interactions (*18, 24*). We firstly identified SE-MYH1/4 as an EDL-specific SE with stronger H3K27ac enrichment and chromatin accessibility in EDL than in SOL. While SE-*fMyh* functions in both mouse SOL and EDL, SE-*MYH1/4* shows EDL-specific activity and drives fast-type fiber specification through chromatin interactions. This highlights the species-specific epigenetic states of SEs controlling fast-type myosin expression.

SE activity depends on TF cooperation and changes in TF enrichment (*41*). Prior studies demonstrated that *Myog* and *Myod* bind to the promoter of *Myh7*, enhancing its transcriptional activity in mice (*24*). Consistent with this, our result revealed that the TF binding sites for *Myog* and *Myod* are detected in SE-MYH7 in pig SOL. Notably, we validated that KLF5 can bind at SE-MYH1/4 to promote its enhancers activity and frequency of interaction with the fast-type *Myh* genes. Additionally, not common with the other KLFs, our results suggested that KLF2 expressed higher in SOL than EDL, and might be activated by a SOL-specific SE. This suggested that KLF2 might be involved in the formation of slow-type fibers. Collectively, these findings establish SE- mediated chromatin interactions and coordinated TF regulation as key determinants of muscle fiber phenotype.

Adult muscle fibers, despite their specified types, can undergo transformation in response to functional demands. These transformations are accompanied by various physiological and biochemical adaptations in skeletal muscle, including mitochondrial biogenesis, angiogenesis, ATP metabolism, and muscle contraction (*42*). Such adaptations are tightly regulated by key genes and gene families associated with muscle development, particularly the well-known MRFs family, Pax family, and the MSTN gene. Additionally, signaling pathways such as the FoxO signaling pathway, PGC-1α pathway, Wingless/Integrated (Wnt), and Sonic Hedgehog (Shh) signaling pathways play critical roles in facilitating muscle fiber-type transformations (*43*).

In this study, we demonstrated for the first time that *STARD7* can drive the transformation of fast-type fibers to slow-type fibers both *in vivo* and *in vitro*. Using CRISPR/Cas9 genome editing, we showed that *in situ* deletion of SE-*STARD7* effectively prevents the expression of the *STARD7* gene. Previous studies have established individual relationships between *STARD7* and intracellular ROS levels (*32*), ROS level and MAPK signaling (*33, 34*), and MAPK signaling and muscle fiber-type transformation (*35*). However, the coordinated regulation of these factors in muscle development had not been explored. Our findings validate that *STARD7* promotes fast-to-slow fiber transformation by reducing ROS levels and inhibiting ERK MAPK signaling activity. In summary, we identified and characterized a previously unrecognized SE that activates *STARD7* expression, elucidating its function and regulatory mechanism in muscle fiber-type transformation.

Muscle is a heterogeneous tissue, with over 68% myonuclei originating from myofibers (*44*). The epigenetics status of CREs and 3D chromatin structure in purified-myonuclei is crucial for investigate muscle fiber-type development, but remains poorly understand due to small sample sizes and complex isolation procedures. Previous studies reported that the trend of chromatin accessibility differences of enhancers between purified myonuclei of slow-type and fast-type muscles is coordinated with the H3K27ac signal between bulk tissues (*18*). In this study, we also revealed a high similarity of H3K27ac signal of myonuclei and bulk tissue of SOL and EDL muscles. These results suggest that it would be feasible to use bulk tissue to reflect the information of epigenetics and 3D chromatin organization of purified-myonuclei. Thus, our multi-omics datasets can provide valuable resources to study the coordination regulation of CREs and distal chromatin looping to shape distinct muscle fibers.

In conclusion, our study provides critical insights into the principles of 3D genome organization as they relate to epigenetic regulation of gene expression in distinct muscle types. We demonstrated that chromatin rewiring at multiple genomic scales and the epigenetic states of CREs underlie transcriptome reprogramming, ultimately dictating the contractile and metabolic properties of specialized muscles. Furthermore, we uncovered the pivotal role of SEs in muscle fiber-type specification and validated KLF5 as a key transcription factor driving the specification of myotubes into fast-type fibers. Lastly, we established that *STARD7* is activated by a distal SE, enabling its role in inducing the transformation of fast-type fibers into slow-type fibers through ROS reduction and ERK MAPK signaling inhibition.

## Materials and Methods

### Ethics Statement

Animal maintenance and experimental procedures were conducted according to Regulations for the Administration of Affairs Concerning Experimental Animals (Ministry of Science and Technology, China, revised in March 2017) and approved by the animal ethical and welfare committee of South China Agricultural University (Permit Number 2021F036, Permit Date 2 March 2021).

### Animals and Sample Collection

Three 6-month-old female Duroc pigs were obtained from the breeding pig farm of Guangdong Wen’s Foodstuffs Group Co., Ltd. (Yunfu, China). These animals were allowed access to feed and water ad libitum and were humanely killed as necessary to ameliorate suffering. Muscle samples from different anatomical sites were rapidly manually separated from each carcass, immediately flash-frozen in liquid nitrogen, and stored at −80℃ or fixed in 4% paraformaldehyde for histology staining.

### RNA-seq library construction and sequencing

The total RNA of tissues or cells was extracted using TRIzol reagent (Invitrogen, 15596026). RNA integrity was qualified by Bioanalyzer 2100 (Agilent, USA) and confirmed by agarose gel electrophoresis. Extracted RNA was used to construct strand-specific libraries with NEBNext® Ultra™ Directional RNA Library Prep Kit (NEB, USA). The libraries were sequenced on the Illumina Novaseq™ 6000 (PE150) platform.

### ChIP-seq library construction and sequencing

Approximately 25 mg of muscle tissues were minced with a scalpel, washed in cold PBS buffer, cross-linked by formaldehyde, and quenched with Glycine Solution. Afterward, the samples were lysed and sonicated to obtain soluble sheared chromatin. Half of the soluble chromatin was collected as input control at –20 °C for DNA sequencing, and the remaining was used for immunoprecipitation reacting with anti-H3K4me3 (Millipore, 07-473) and anti-H3K27ac (Millipore, 07-360). Chromatin was de-cross-linked, purified, and used to generate libraries. For both input DNA and immunoprecipitated DNA, each library was sequenced on the Illumina HiSeq X Ten (PE150) platform.

### CUT&Tag library construction and sequencing

The muscle tissue was placed into a pre-chilled Dounce homogenizer containing cold lysis buffer. The tissue was homogenized using the A pestle until no resistance was felt, indicating thorough disruption of the tissue. The homogenate was then pre-cleared by filtration through a 100μm nylon mesh filter to remove debris. Subsequently, the filtered homogenate was further homogenized using the B pestle for 20 strokes to ensure complete lysis. The supernatant was carefully transferred to a low-binding tube. Nuclei were pelleted by density gradient centrifugation at 3000g for 30 minutes in a solution composed of 60% Percoll and 2.5M sucrose. The collected nuclei were incubated with concanavalin A-coated magnetic beads. Cell-bound beads were resuspended and then immunoprecipitated with anti-H3K27me3 (Millipore, 07-449) and anti-CTCF (Millipore, 07-729) with rotation at 4 °C overnight. The enzyme pA-Tn5 Transposase precisely binds the DNA sequence near the target protein under the antibody guidance. The DNA sequence is tagmented, with adapters added at the same time at both ends, which can be enriched by PCR to form the sequencing-ready libraries. After the PCR reaction, libraries were purified with the AMPure beads (Beckman, A63881), and library quality was assessed on the Agilent Bioanalyzer 2100 system. The constructed library was sequenced on the Illumina Novaseq™ 6000 (PE150) platform.

### ATAC library construction and sequencing

Approximately 5 mg of tissue was cut into small pieces and crushed into a fine powder in liquid nitrogen. The pulverized tissues were resuspended in lysis buffer (50 mM HEPES with pH 7.5 (Life Technologies, 15630-080), 140 mM NaCl (Ambion, AM9760G), 1 mM EDTA with pH 8.0 (Ambion, AM9260G), 10% glycerol (Sigma, G7757), 0.5% NP-40 (Roche, 11754599001), and 0.25% Triton X-100 (Amresco, 0694), followed by isolation of 50,000 nuclei using a previously published protocol with minor modifications (*45*). Then, the isolated nuclei were incubated in the transposition reaction mix (12.5 μL TD buffer, 10 μL ddH_2_O, and 2.5 μL TDE (Illumina, FC-121-1030) at 37 °C for 1 h, and then purified with a QIAGEN MinElute PCR Purification Kit (QIAGEN, 28006). Next, an average of 11 cycles of PCR was performed with transposed DNA and amplified libraries were run on an Agilent TapeStation 2200 (Agilent Technologies, USA) using a D5000 DNA ScreenTape to assess their quality via visualizing nucleosomal laddering. The final libraries were quantified and sequenced on the Illumina HiSeq X Ten (PE150) platform.

### BL-Hi-C library construction and sequencing

Approximately 5 × 10^6^ isolated cells were resuspended in 1 ml of 1% SDS lysis buffer (50 mM HEPES-KOH, 150 mM NaCl, 1 mM EDTA, 1% Triton X-100, 1% SDS) at room temperature for 15 minutes and centrifuged at 500 × g for 2 minutes. The pellet was washed with 1 ml 0.1% SDS lysis buffer and centrifuged at 500 × g for 2 minutes. The supernatant was discarded, and the cells were incubated with 50 µl of 0.5% SDS for 30 minutes at 37 °C. The SDS was quenched by the addition of 145 µl H_2_O and 12.5 µl 20% Triton X-100, and the genome was digested overnight at 37 ℃ with 100U AluI (NEB, R0137V) into blunt-end fragments. Cleaved chromatin was A-tailed by adding 10 mM dATP solution (NEB, N0440S) and Klenow Fragment (3′→5′ exo-) (NEB, M0212L) with rotation at 37°C for 40 minutes. Then, chromatin was treated with adenine and ligated with a biotinylated bridge linker (F: CGCGATATC/bio-T/TATCTGACT; R: GTCAGATAAGATATCGCGT) at 16°C for 4 h. After ligation, the nuclei with in situ proximity ligation were treated with l-exonuclease (NEB, M0262S) and exonuclease I (NEB, M0293V) for 1h at 37℃ to remove noise DNA contain no ligation DNA or bridge linker. The DNA was obtained by proteinase K digestion at 55°C overnight and purified by phenol-chloroform extraction with ethanol precipitation. Then, the DNA was fragmented into 300 bp on average by sonication, and the biotin-labeled DNA fragments were pulled down with streptavidin-coated M280 beads. The DNA library was sequenced using the BGISEQ DNBSEQ-T7 (PE150) platform (BGI lnc., Shenzhen, China).

### RNA-seq data analysis

Clean reads were aligned to the pig reference genome with chain-specific parameters: “rna-strandness RF” by Hisat2 (*46*). The gene counts were calculated by featurecounts (*47*) and then normalized to transcripts per kilobase million (TPM). Only genes with TPM ≥ 0.5 in at least one sample were considered as expressed genes and used for further analysis. Differentially expressed genes were identified with FDR < 0.05 and fold change > 1 using DESeq2 (*48*). GO analysis was assessed using PANTHER, and GSEA was implemented by the R package Clusterprofiler (*49, 50*). KEGG pathway analyses were carried out in KOBAS-I (*51*).

### ChIP-seq and CUT&Tag data analysis

Clean reads were mapped to the pig reference genome (susScr11) or mouse reference genome (mm39) using BWA (*52*). Picard was used to obtain unique mapped reads without PCR duplicates (http://broadinstitute.github.io/picard/). For the H3K27me3 dataset, peaks were called using MACS2 (*53*) with parameters (-g 2.8e+9 -q 0.05 --keep-dup all --nomodel –broad). For H3K4me3, H3K27ac, and CTCF datasets, peaks were called using MACS2 with parameters (-g 2.8e9 -p 0.00001 --shift 0 --extsize 150 --nomodel -B --SPMR --keep-dup all --call-summits). Peaks were further processed by following four steps as previous study (*54*): (i) narrow peaks with *P* < 10^−5^ were remained; (ii) read coverage of 2 kb region centered at the midpoint of peaks was calculated by Deeptools (*55*) and normalized with read depth (IP_RPM_ = IP _peak region total reads_ / IP _total mapped reads per million_; INPUT_RPM_ = INPUT _peak region total reads_ / INPUT _total mapped reads per million_); (iii) the enriched regions for H3K4me3 and H3K27ac were defined with more than a two-fold change of normalized read coverage (IP_RPM_ > 2 × INPUT_RPM_) and with normalized read coverage change (IP_RPM_-INPUT_RPM_) > 1; (iv) the overlap of enriched regions were merged, and then the 2kb regions centered at the midpoint of merged regions were acquired to identify enhancers and promoters. Motif analysis was performed using HOMER’s findMotifsGenomDe.pl program with parameters (-len 8,10,12 -gc) (*56*).

### ATAC-seq data analysis

ATAC-seq datasets were processed with reference to the ENCODE ATAC-seq pipeline (https://github.com/kundajelab/atac_dnase_pipelines). After checking and trimming the adapter with cutadapt (*57*), the clean reads were mapped to the pig genome (susScr11) using Bowtie2 with parameters (--very-sensitive -X 2000) (*58*). The reads with low MAPQ reads (<25), reads mapped to mitochondrial chromosome, and PCR duplicated reads were removed using samtools (*59*) and Picard. The peak for each sample was called individually with MACS2 (-g 2.8e+9, -p 0.01, --nomodel, --shift - 75, –extsize 150, -B, --SPMR, --keep-dup all, --call-summits). The significant peaks (*P* < 10^−5^) of each sample were used for further analysis. Then, the filtered peaks of replicates were merged with BEDtools (at least 50% overlap between two replicates).

### Chromatin state analysis and characterization of CREs

ChromHMM (version 1.22) was used to learn and characterize chromatin states (*27*). Candidate enhancers were defined as H3K27ac peaks not overlapping with H3K4me3 peaks or TSS regions (3.0 kb upstream to 1.5 kb downstream of TSS) overlapped with S3 or S4 regions. Candidate promoters were defined as H3K4me3 peaks overlapped with S5 or S6 regions. Candidate silencers were defined as H3K27me3 peaks overlapped with S1 regions. SEs were identified using the ROSE algorithm with default parameters, and all enhancers were used as input constituent enhancers. The CREs with no overlap in other tissues were defined as tissue-specific CREs. Candidate CREs (promoters and enhancers) were converted from pig genomic (susScr11) to human/mouse genomic locations (hg19/mm9) by using the LiftOver tool with the parameter “minMatch = 0.5”. The CRE that could be converted to human/mouse genomic locations was considered as sequence-conserved CREs. Furthermore, the CREs were considered usage-conserved if the corresponding homologous sequence was annotated as human/mouse CREs in the FANTOM5 database (*60*).

### Footprint analysis

The vertebrates TF motifs were downloaded from JASPAR (2024) (*61*). The binding score of each TFs and differential TF binding analysis were performed by TOBIAS with default parameters (*62*). Only TFs with TPM ≥ 0.5 in at least one sample were considered as expressed TFs and used for further analysis.

### BL-Hi-C data analysis

The adapter and linker sequence of raw Hi-C reads was trimmed using cutadapt (*57*). These processed clean data were used to generate contact matrices using the HiC-Pro pipeline (*63*). Chromatin A/B Compartment was identified using cooltools at 50kb resolution (*64*). TAD and TAD boundaries were identified using HICExplorer at 50kb resolution (*65*). The definition of cTADs and their intra-TAD interaction strength (D-score) was calculated as reported in the previous study (*66*). Loop calling was performed by HICCUPS in 5kb, 10kb, and 25kb resolution (*67*). The aggregate enrichment of TADs and loops was generated by coolpup (*68*). Genome-wide interaction and heatmap were visualized using the WashU EpiGenome browser (*69*). Genomeflow is set 3DMax model to construct 3D chromatin conformation model with sparse contact matrix as input at 10kb resolution (*70*). The contact frequency of loops was calculated by hicPlotViewpoint function in HICExplorer. The interactions were then normalized by the number of interactions in the whole chromosome, and used to generate virtual 4C profiles.

### Isolation of PSCs

For PSCs isolation, hindlimb muscles of newborn piglets within one week were cut into pieces and digested with 2mg/ml type II collagenase (Sigma, 9001-12-1) at 37 °C for 2.5 h. The tissue suspension was then filtered through 100-, 200-, and 400-mesh sieves and washed in DPBS (Gibco, 14190144) with 1% penicillin–streptomycin (Gibco, 15140122). Single cells were collected by centrifugation at 1800 × g. Subsequently, cells were differentially adhered for 2 times to obtain purified PSCs.

### Cell culture

For proliferation, PSCs were cultured in RPMI-1640 medium (Gibco, 11875119) with 20% FBS (Gibco, 10099141), 1% non-essential amino acids (Gibco, 11140050), 0.5% chicken embryo extract (GEMINI, 0928501), 1% GlutaMax (Gibco, 35050061), 1% antibiotic–antimycotic (Gibco, 15140122), and 2.5 ng/ml bFGF (Gibco, 13256029) under moist air with 5% CO_2_ at 37°C. For differentiation, the cells that reached 80-90% confluence were switched to differentiation in RPMI 1640 medium containing 2% horse serum (HyClone, SH30074.03).

### Cell transfection

All small interfering RNA (siRNA) sequences used in this study are presented in table S9. Lipofectamine 3000 (Invitrogen, L3000150) and jetPRIME reagent (Polyplus transfection, 114-15) were respectively used for the transfection of plasmids and siRNA according to the instructions.

### Quantitative real-time PCR (qPCR)

Total RNA was reverse-transcribed to create cDNA by Evo M-MLV RT Kit (Accurate Biotechnology, AG11711) following the manufacturer’s manual. The qPCR reaction was performed on Quant Studio 7 Flex Real-Time PCR System (Thermo Fisher Scientific, USA). The relative expression level of one gene was determined by the 2^−ΔΔCt^ method. All primers used in this study were presented in table S9.

### Chromatin Immunoprecipitation qPCR (ChIP-qPCR)

Chromatin immunoprecipitation experiment was carried out by using ChIP kit (Beyotime, P2078) according to the recommended protocol. Briefly, 4 × 10^6^ PSCs were cultured to 90% confluence and fixed for 10 min at room template with 1% formaldehyde. The crosslinking was stopped by adding 125 mM glycine. Cell lysates were sheared by sonication in 1% SDS lysis buffer to generate chromatin fragments, followed in sequence with immunoprecipitation, reversal of cross-links, and DNA purification. Two micrograms of anti-KLF5 (Active Motif, 61099) and H3K27ac anti-H3K27ac (Millipore, 07-360) or negative control IgG was used to each immunoprecipitated reaction. Fold enrichment was quantified using RT-qPCR. All primers used in this study were presented in table S9.

### Western Blotting (WB)

Tissue or cells were lysed in RIPA buffer (Beyotime Biotechnology, P0013C) with 1% (v/v) PMSF (Beyotime Biotechnology, ST505). Denatured protein samples were separated by SDS-PAGE and transferred to the 0.45 um PVDF membranes (Millipore, IPVH00010). After blocking with 5% non-fat milk, the proteins in membranes were subjected to immunoblotting analysis with primary and secondary antibodies. The membranes were developed with an ECL Kit (Millipore, WBULS0500) for visualization. The antibodies used in this study included anti-MyhC IIb (ProteinTech, 20140-1-AP), anti-MyhC I (ProteinTech, 22280-1-AP), anti-STARD7 (ProteinTech, 15689-1-AP), anti-ERK1/2 MAPK (ProteinTech, 11257-1-AP), anit-phosphop-ERK1/2 MAPK (Proteintech, 28733-1-AP), anti-β-tubulin (Servicebio, GB11017), anti-GAPDH (Absin, abs132004), and secondary antibodies goat anti-rabbit IgG (Beyotime, A0208). All protein levels were normalized to corresponding β-tubulin or GAPDH, and densitometric quantification of the bands was performed using ImageJ software.

### Immunofluorescence staining

Cell immunofluorescence staining was performed according to a previously published method (*71*). Multiplex immunofluorescence staining based on tyramide signal amplification was performed by using four-color multiple fluorescent immunohistochemical staining kit (Absin, abs50012) according to the manufacturer’s protocols. Immunofluorescence staining antibodies included anti-Fast Myosin Skeletal Heavy chain (Serviebio, GB112130-100), anti-Slow Skeletal Myosin Heavy chain (Serviebio, GB112131-100), anti-KLF5 (Active motif, 61099) and secondary antibodies biotinylated goat anti-mouse IgG (BIOSS, bs-0296G-Bio), and Alexa Fluor 488-labeled goat anti-rabbit IgG (H + L) (Beyotime, A0423). Images of at least three random fields of view were captured using a Nikon ECLIPSE Ti microscope (Nikon, Tokyo, Japan) and quantified using ImageJ.

### CRISPR-cas9 experiment

The CRISPR/Cas9 plasmid pX330-U6-Chimeric_BB-CBh-hSpCas9 (pX330) was obtained from Addgene (Plasmid #42230). The sgRNAs against the selected enhancers were listed in table S9. The sgRNAs were designed, synthesized, annealed, and cloned into px330 plasmid to form the recombinant plasmids. After the confluence of PSCs reached 80-90%, we transfected the recombinant pX330-sgRNA plasmids, together with pcDNA3.1(+) plasmids (containing neomycin resistance) to cells by lipofectamine 3000 (Invitrogen, L3000150). After 36 h, G418 (100 μg/mL; Sigma, G8168) selection of the cells was performed for approximately 2 weeks. Then, the mixed population of cells was collected to extract RNA and DNA. The genomic DNA of collected cells was isolated according to the protocol of the MicroElute Genomic DNA Kit (OMEGA, D3096) for PCR. PCR was performed using the specific primers (table S9). The deletion efficiency was analyzed by 1% agarose gel electrophoresis and sanger sequencing of PCR product.

### Validation of loops by 3C-qPCR

The isolated muscle cells from muscles tissues or PSCs were cross-linked in 1% formaldehyde for 10 minutes at room temperature and then quenched by adding glycine (final to 0.125 M). The samples were recovered by centrifugation and washed with 1 mL DPBS (Gibco, 14190144). Each library was cleaved with 0.2% NP40 (Sigma, 74385), transferred to the enzyme digestion reaction system, and incubated with 100U Mbol restriction enzyme (NEB, R0147S) at 37 ℃ for 2 h. Centrifuge at 3000 g at 4 ℃ for 10 minutes and discard the supernatant. For ligation, T4 DNA ligase (NEB, M0202M) was treated for 5h at 16 °C and for 2 h at room temperature. Ligated DNA was purified by phenol/chloroform/isoamyl-alcohol (25:24:1) extraction and ethanol precipitation. qPCR was performed to calculate the relative interaction frequencies. The 3C-qPCR primers were designed using unidirectional strategy (table S9).

### Detection of intracellular ROS level

The ROS levels after overexpression of *STARD7* in PSCs were detected using ROS assay kits (Beyotime, S0033M). Briefly, the PSCs were harvested and added to DCFH-DA solution diluted with serum-free medium. The cell suspension was incubated with the DCFH-DA for 30 min with shaking every 5 min in the dark. At the end of the incubation period, the fluorescence intensity was detected by using BioTek’s Synergy™ HT Reader.

### Cell metabolism assays

Seanorse XF Glycolytic Rate Assay Kit (Seahorse, 103344-100) was used to detect the extracellular acidification rate (ECAR) reflected the glycolysis capacity of PSCs. Briefly, cells (4 × 10^4 per well) were seeded and cultured in Seahorse XFe24 cell plate (Seahorse, 102340-100) before conducting the assay. After transfection, the media was changed to Seahorse XF DMEM Medium (pH 7.4, Seahorse, 103575-100) supplemented with 10 mM glucose (Seahorse, 103577-100), 1 mM pyruvate (Seahorse, 103578-100) and 2 mM glutamine (Seahorse, 103579-100). After cultured at 37°C in a CO2-free incubator for 60 min, the real-time ECAR of cells was first recorded under basal condition as data on basic glycolysis. Through the sequential addition of 0.5 μM rotenone/antimycin A (Rot/AA) followed by 50 mM of 2-depxy-d-glucose (2-DG), the ECAR was recorded and compensatory glycolysis were calculated by using a Seahorse XFe24 Extracellular Analyzer according to the manufacturer’s protocols.

### Dual-Luciferase reporter assay

The pig genomic DNA was used as a template to amplify enhancers. The amplified products were separated on a 1.5% agarose gel, purified, and then cloned into the modified pGL3-Promoter plasmids. Constructed plasmids were transfected to PSCs using Lipofectamine 3000, and incubated for 24h with three technical replicates for each construct. Luciferase activity was measured by Dual-Luciferase Assay (YEASEN, 11402ES60) according to the manufacturer’s manual, and Renilla luciferase activity was normalized against Firefly luciferase activity.

### Knockdown of STARD7 in vivo by lentivirus infection

Six-week C57 male mice were injected with 100 μL final volumes of lentivirus containing small interfere STARD7 (LV3-shSTARD7) or control (LV3-shNC) at 2 × 10^7^ TU/ml into the right and left GAS muscles of the hind legs, respectively. LV3-shSTARD7 and LV3-shNC were synthesized by GenePharma (China, Shanghai). The mice were injected weekly and killed after 8-weeks injection. GAS muscles were collected once a week. The meat color, including L*(lightness), a*(redness), and b* (yellowness), was measured by CR-400 Chroma meter (Konica Minolta Sensing Ins., Japan) at 45 min postmortem as a mean of three random readings calibrated against a standard white plate (8-mm diameter aperture, d/0 illuminant).

## Acknowledgments

[Dum]

## Author contributions

Conceptualization: B.T., T.G., and Z.W.

Methodology: B.T., and L.X.

Investigation: B.T., L.H., Z.L., G.C., W.S., L.X., L.L., J.W., and G.L.

Bioinformatic analysis: B.T.

Visualization: B.T.

Supervision: E.Z.

Writing—original draft: B.T.

Writing—review and editing: B.T., T.G., Z.W., and L.X.

## Data and materials availability

Sequence data of pig SOL and EDL muscles in this study have been deposited in the NCBI Sequence Read Archive (SRA) under the accession number PRJNA1036842 and PRJNA810786. The RNA-seq data of STARD7 overexpression have been deposited in the SRA under accession number PRJNA1112571. The ChIP-seq and ATAC data of mouse SOL and EDL muscle were downloaded from Gene Expression Omnibus (GEO) under accession number GSE123879. The RNA-seq data have been deposited in PRJNA1112571. The H3K27ac ChIP-seq datasets of purified-myonuclei and bulk tissues are deposited in GSE123879 and GSE182667. The TF, histone, ATAC-seq datasets of C2C12 myotubes were download from PRJNA305993, PRJNA844370, PRJNA320053, PRJNA260504, PRJNA185319, PRJNA63475, PRJNA268849, PRJNA734569, and PRJNA1081841. All other data needed to evaluate the conclusions are present in the paper and/or the Supplementary Materials. Source data are provided with this paper.

## Code availability

Custom scripts described in the Methods will be made available upon request.

## Supplementary Figures

**Fig. S1.**
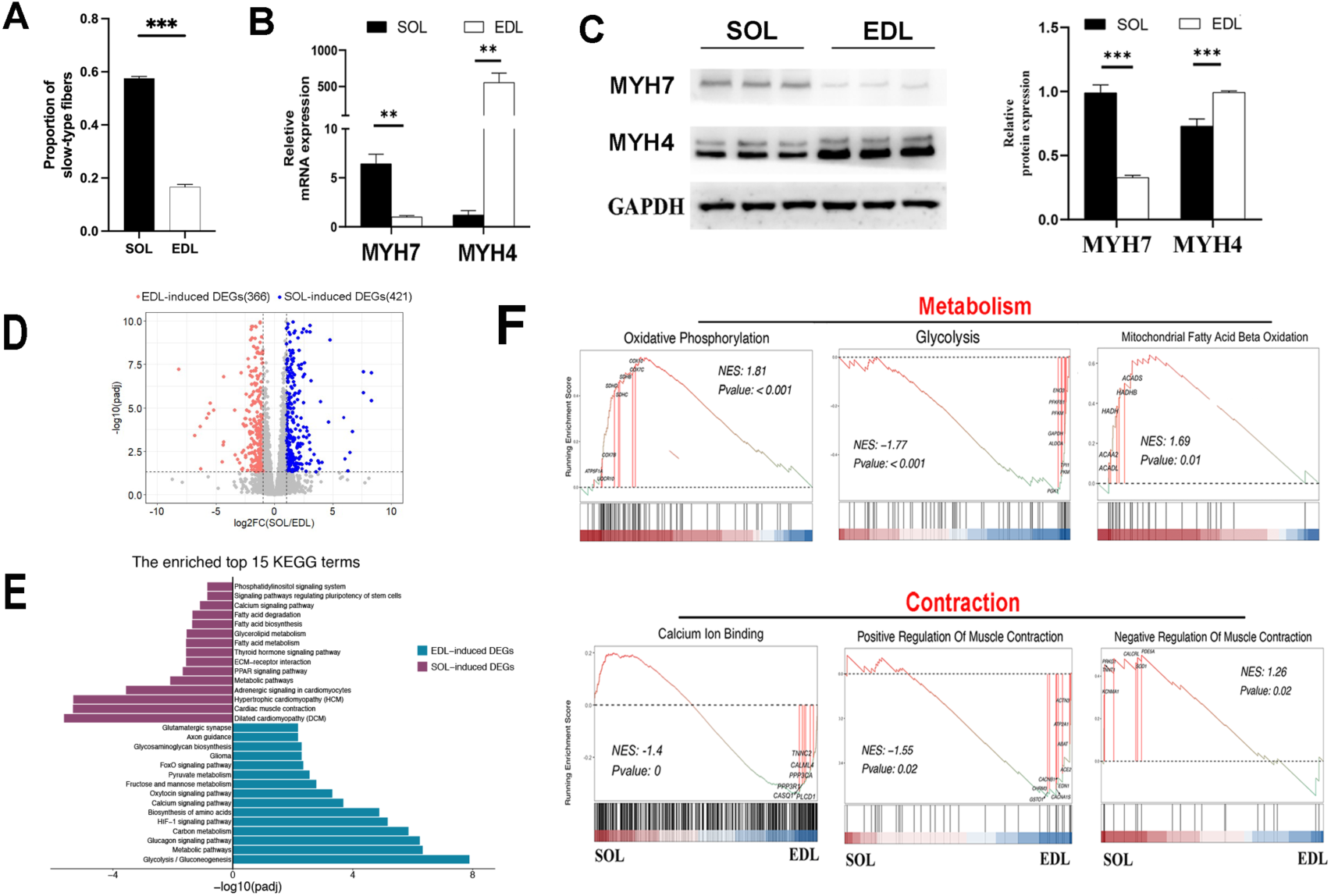
Phenotypic differences and transcriptome dynamics in SOL and EDL. (A) Statistical analysis showing the changes in the proportions of fast-type and slow-type fibers. (B-C) qPCR and Western blot analyses revealing differences in the expression of MYH4 and MYH7 between SOL and EDL. (D) The volcano plot of DEGs identified by RNA-seq. SOL-induced DEGs: DEGs that highly expressed in SOL; EDL-induced DEGs: DEGs that highly expressed in EDL. (E) KEGG pathway enrichment analysis of SOL-induced DEGs or EDL-induced DEGs. (F) GSEA plot highlighting the significant enrichment of selected pathways in SOL and EDL. Genes within each pathway are ranked by log2(fold change in gene expression) (SOL/EDL), with blue indicating genes upregulated in EDL and red indicating those upregulated in SOL. NES: Normalized Enrichment Score.

**Fig. S2.**
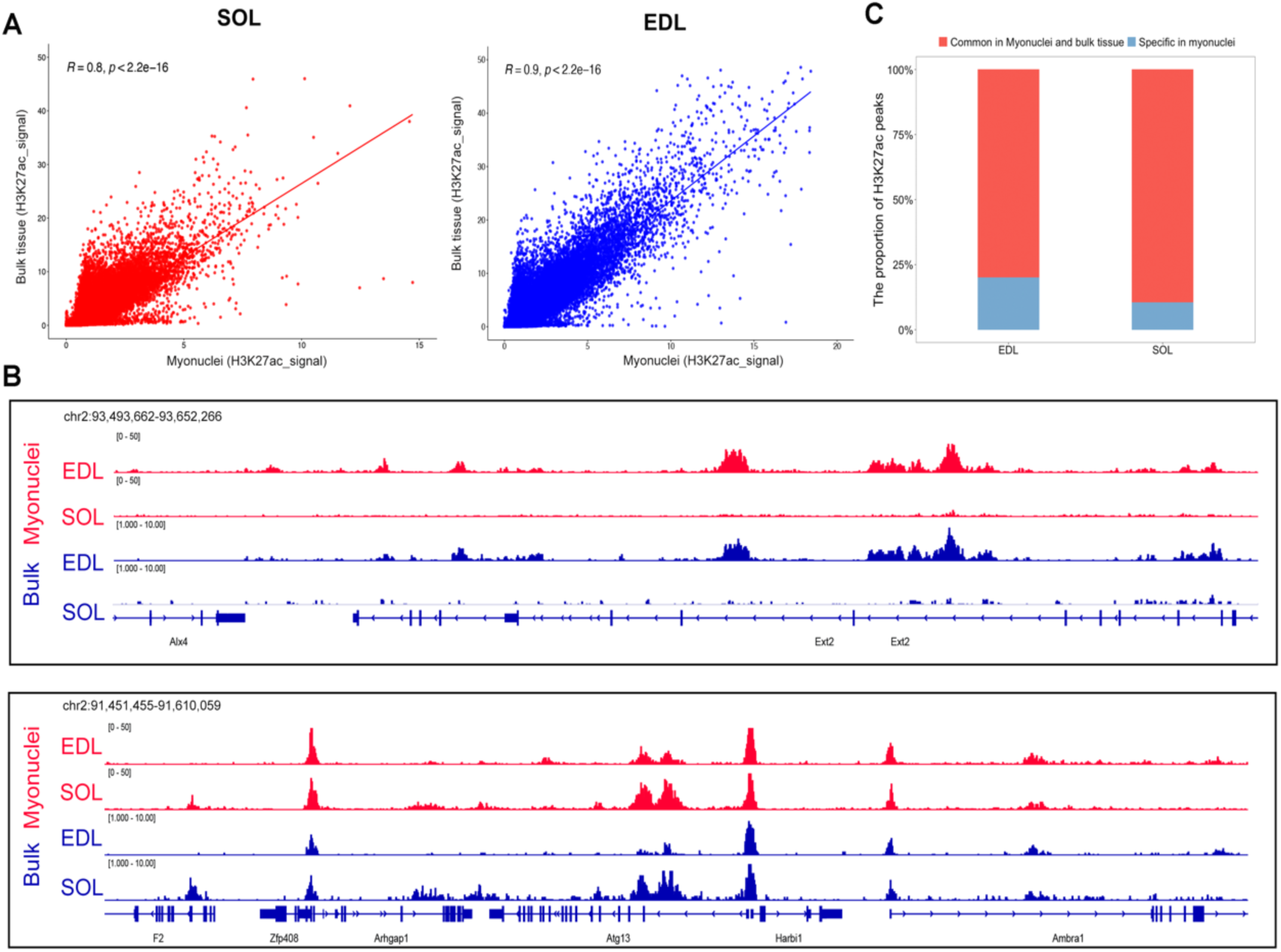
The similarity of H3K27ac signal of bulk tissues and purified-myonuclei of mouse SOL and EDL muscles. (A) The Pearson Correlation Coefficient of genome-wide H3K27 signal in the bulk tissue and myonuclei of SOL and EDL muscles. (B) The ChIP-seq track of H3K27ac signal of indicated regions. (C) The overlap of peaks identified from myonuclei and bulk tissues.

**Fig. S3.**
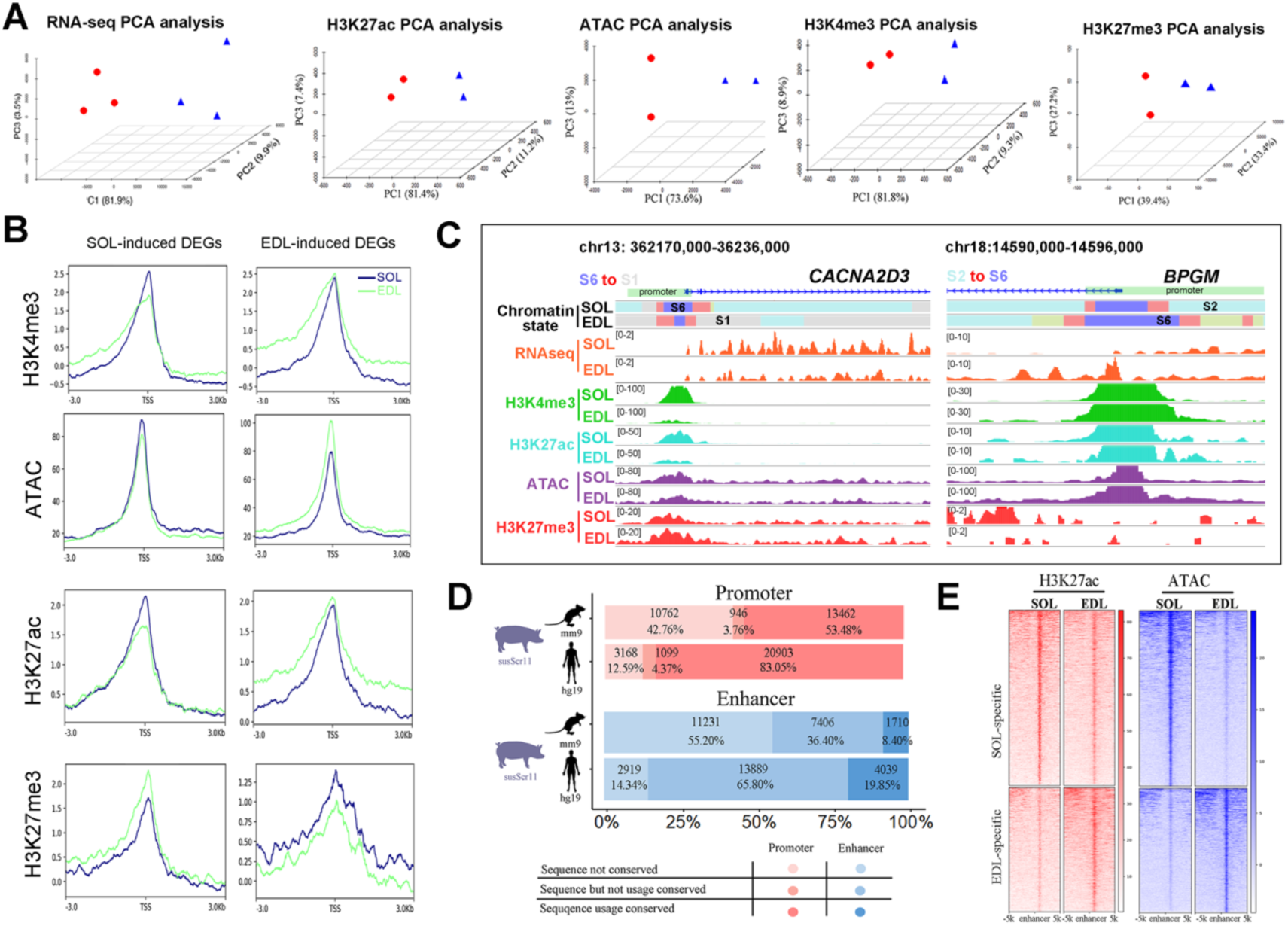
The quality of epigenetics datasets and CREs conservation in SOL and EDL. (A) PCA plot based on RNA-seq, ATAC-seq, H3K27ac/H3K4me3 CUT&Tag, and H3K27me3 ChIP-seq data. (B) Enrichment plot showing that active histone marks (H3K27ac, H3K4me3) and chromatin accessibility are positively correlated with gene expression, while H3K27me3 is negatively correlated. (C) Tracks showing ATAC-seq, chromatin state, and histone modification enrichment levels at CACNA2D3 and BPGM in SOL and EDL. Promoter regions are marked in green. (D) Conservation of sequence and use of CREs across human, mouse, and pig genomes. (E) Enrichment plot showing levels of H3K27ac and chromatin accessibility in tissue-specific enhancers.

**Fig. S4.**
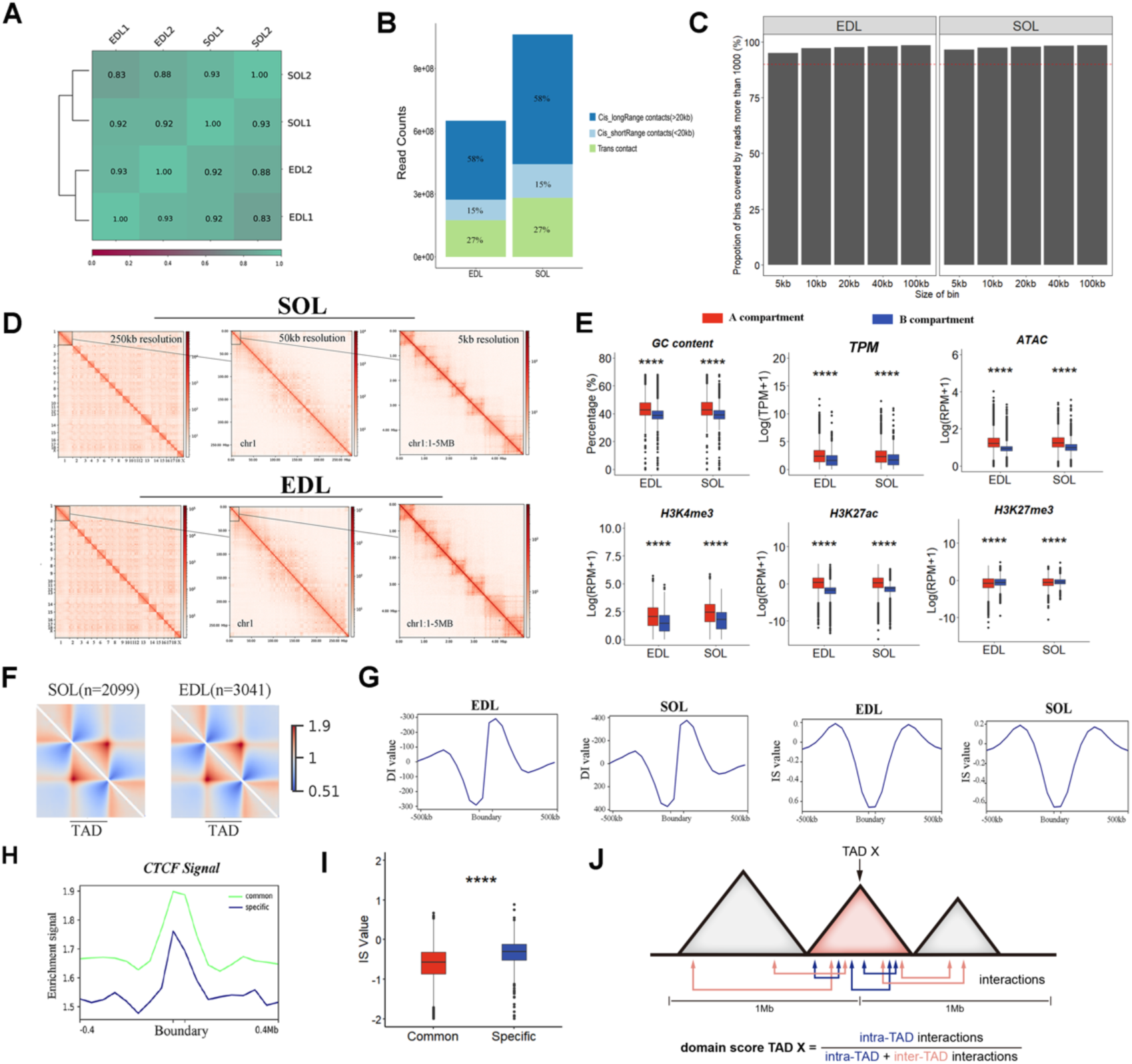
Summary of Hi-C libraries and characteristics of compartments and TADs. (A) Correlation heatmap showing PCCs of genome-wide interactions at 50 kb resolution in SOL and EDL. (B) Distribution of Hi-C valid pairs. (C) Bar plot displaying the proportion of bins with over 1000 reads at specified distances, with the red dotted line indicating 80% of bins containing more than 1000 reads. (D) Chromatin interaction heatmaps for chromosome 1 at 250-kb, 50-kb, and 5-kb resolutions in SOL and EDL. (E) Comparison of GC content, histone modifications, chromatin accessibility, and gene expression between compartments A and B. (F) Aggregate heatmaps representing average *cis*-chromatin contact conformation around identified TADs. (G) Mean values of IS and DI around TAD boundaries. (H-I) Comparison of CTCF binding signal (H) and IS value (I) at common and specific TAD boundaries. (J) Cartoon depicting D-score calculation. Arrows indicate intra- or inter-TAD interactions. Statistical significance: \**P* < 0.05, \*\**P* < 0.01, \*\*\**P* < 0.001, \*\*\*\**P* < 0.0001, two-sided Wilcoxon test.

**Fig. S5.**
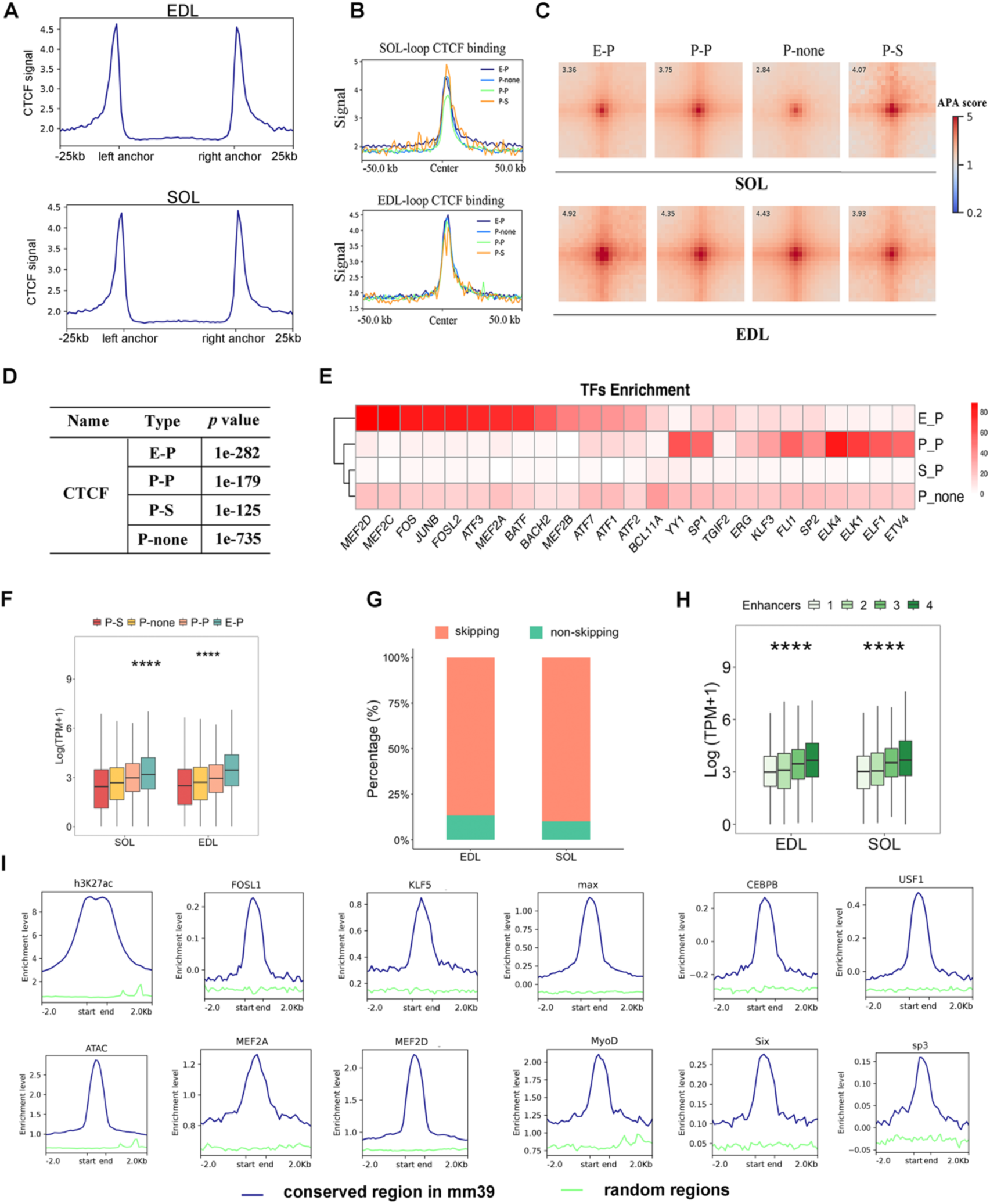
CTCF binding at loop anchors and characteristics of E-P interaction. (A) Enrichment of CTCF at loop anchors. (B-C) Enrichment plot and APA plot showing a positive correlation between CTCF binding at loop anchors and the strength of loop interactions. (D-E) Enriched TFs in classified loops. (F) Gene expression levels of genes associated with different CREs. (G) Percentage of enhancers interacting with distant promoters rather than the closest ones. (H) Gene expression levels of genes that loop with different numbers of enhancers. (I) The TF enrichment plot of identified enhancers regions in C2C12 myotube. \*\*\*\**P* < 0.0001, two-sided Wilcoxon test.

**Fig. S6.**
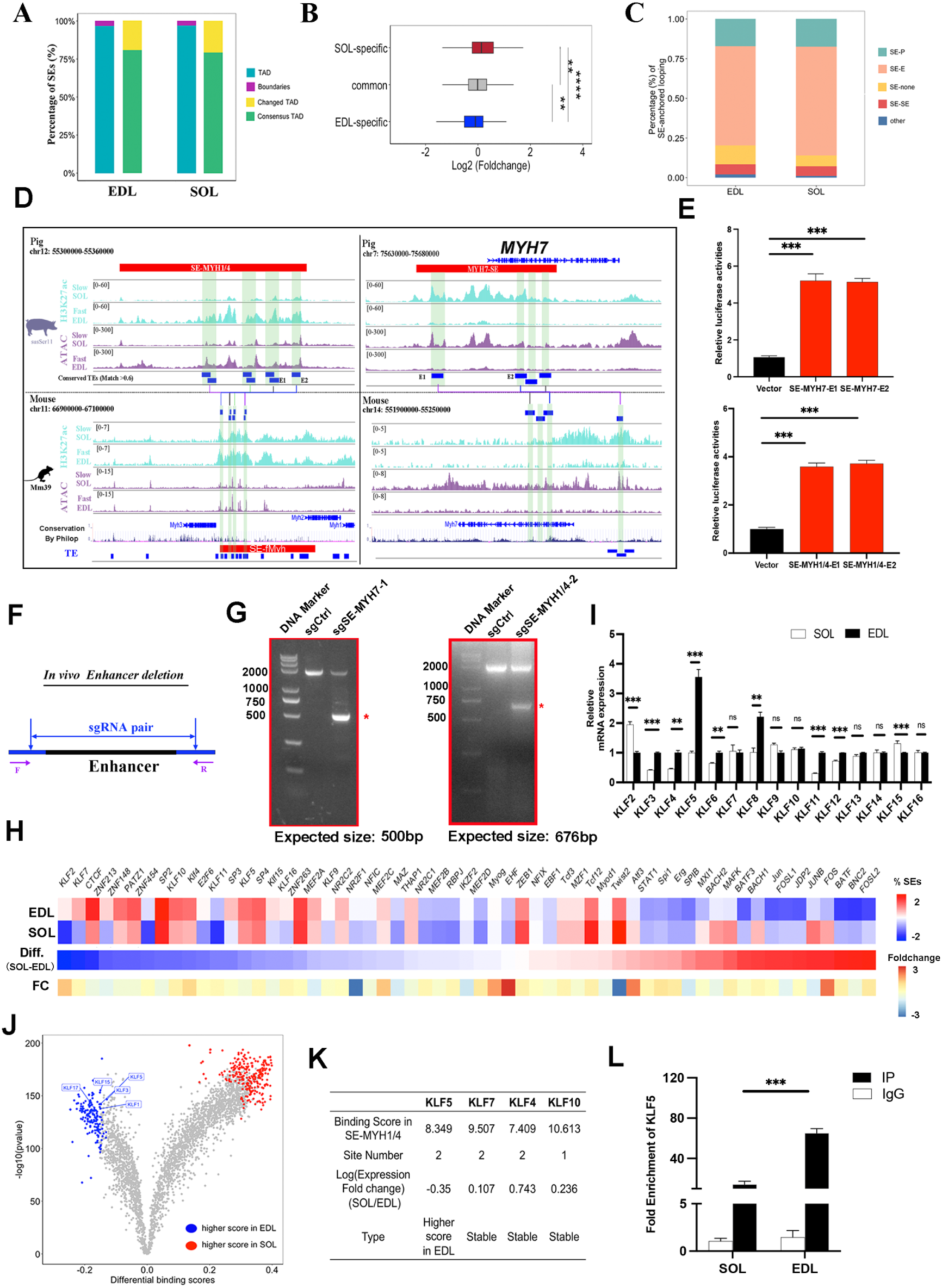
TF enrichment, conservation and validation of SEs. (A) Location of SEs. (B) Expression levels of genes proximal to different SEs. (C) Proportion of SE-anchored loop interactions. (D) The conservation analysis of SE-MYH7 and SE-MYH11/4 between SOL and EDL muscles of pig and mouse. (E) Dual-reporter assay detecting the enhancer activity of SE-MYH7 and SE-MYH1/4 in differentiated PSCs. (F) Schematic illustrating the design of sgRNA pairs for *in vivo* deletion of individual regulatory elements in PSCs via CRISPR-Cas9. Purple arrows (F and R) indicate PCR primers used for genomic PCR analysis of *in vivo* deletion efficiency. (G) DNA isolated from control (sgCtrl) or deletion (sg-SE-MYH1/4-2) PSCs was analyzed by genomic PCR to assess cleavage efficiency. The deletion product (red asterisk) and expected size after deletion are shown. (I) The gene expression of KLFs in SOL and EDL muscles. (H) Enrichment analysis of TFs in SEs. Top panel: Proportion of SEs with at least one TFBS. Middle panel: Proportion differences of SEs with at least one TFBS between SOL and EDL. Diff.: Proportion differences. Bottom panel: Expression changes of TFs between SOL and EDL. FC: Fold change in TF expression (SOL/EDL). (J) Volcano plot indicates the differential binding analysis of TFs. (K) Characterization of KLF transcription factors bound in SE-MYH1/4. (L) ChIP-qPCR showing the enrichment of KLF5 in SE-MYH1/4 in SOL and EDL muscles.

**Fig. S7.**
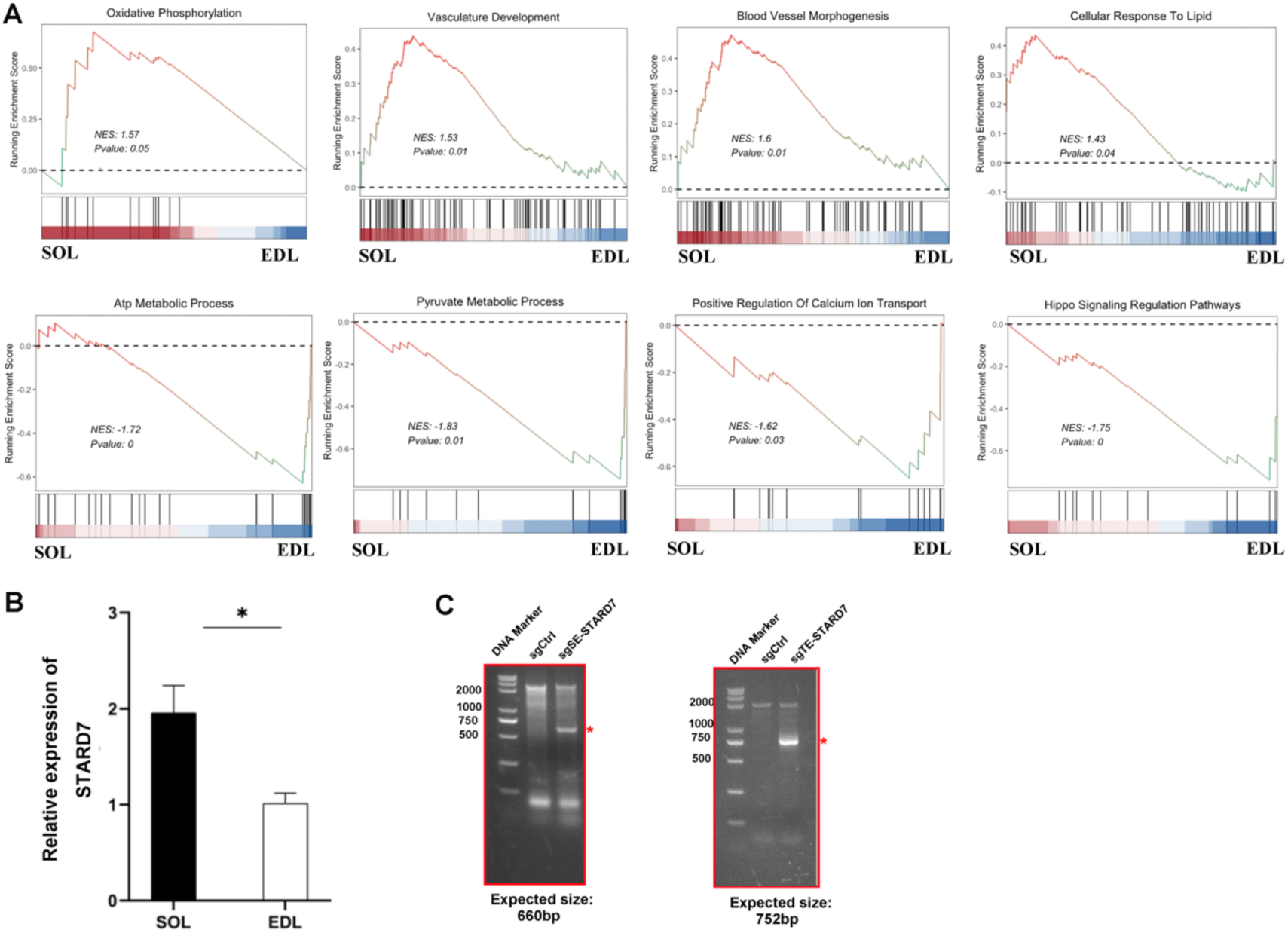
GSEA analysis and SE validation. (A) GSEA plot highlighting the significant enrichment of selected pathways in SOL and EDL. SE-regulated genes within each pathway are ranked by log2(fold change in gene expression) (SOL/EDL), with blue indicating genes upregulated in EDL and red indicating those upregulated in SOL. NES: Normalized Enrichment Score. (B) The expression level of STARD7 in SOL and EDL detected by RT-qPCR. (C) DNA isolated from control or deletion PSCs was analyzed by genomic PCR to assess cleavage efficiency. The deletion product (red asterisk) and expected size after deletion are shown.

**Fig. S8.**
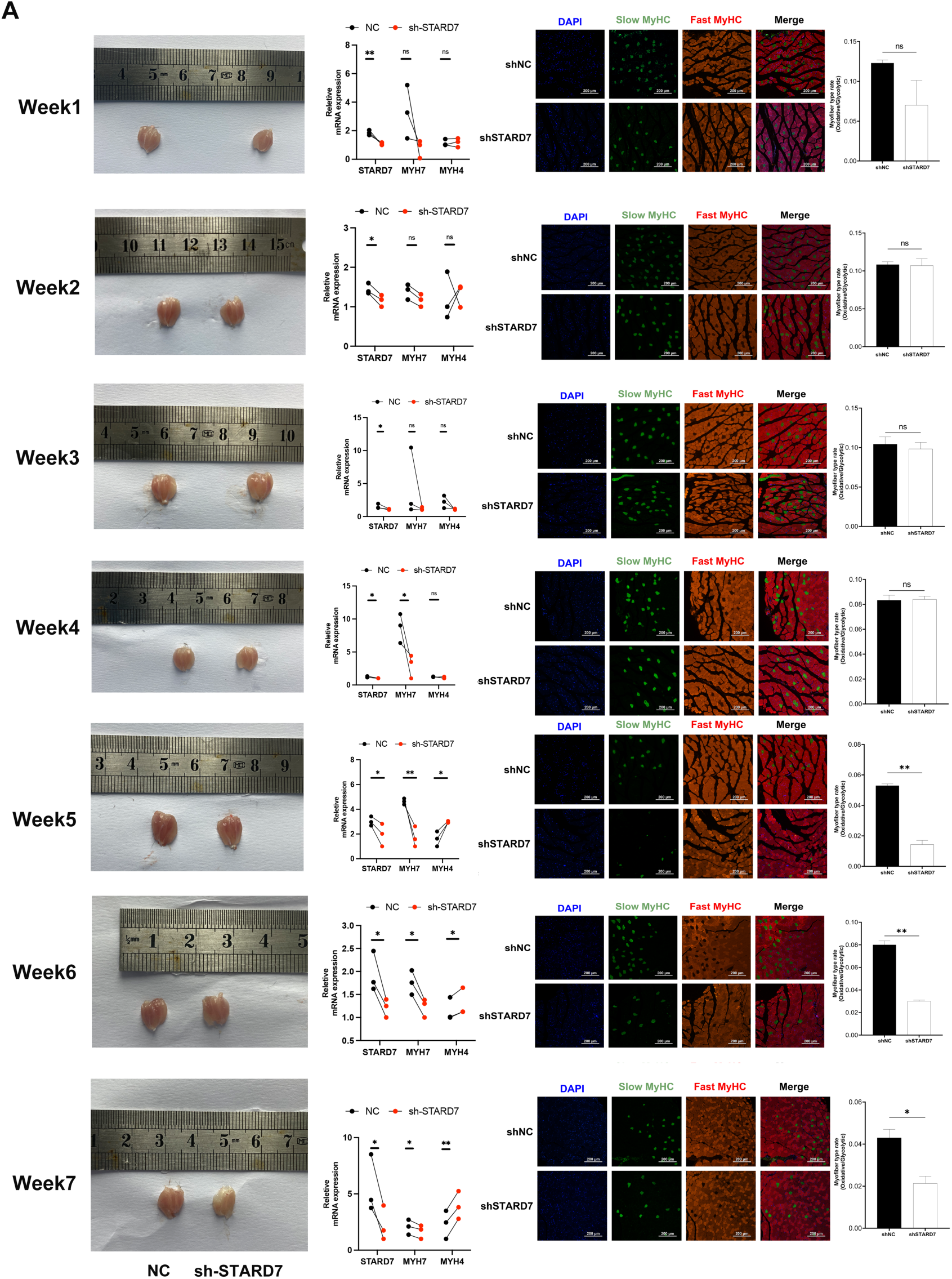
The monitor of fiber-type switching in vivo after STARD7 knockdown in vivo. (A) Left panel: Representative images of the GAS muscles from the left LV3-shNC (NC) or right LV3-shSTARD7(sh-STARD7) muscles (left panel). Middle panel: qPCR depicting gene expression change between NC and sh-STARD7 particles. Right panel: Representative image of immunofluorescence staining and statistical analysis of the proportion changes of fast-type fibers (green) and slow-type fibers (red).

**Fig. S9.**
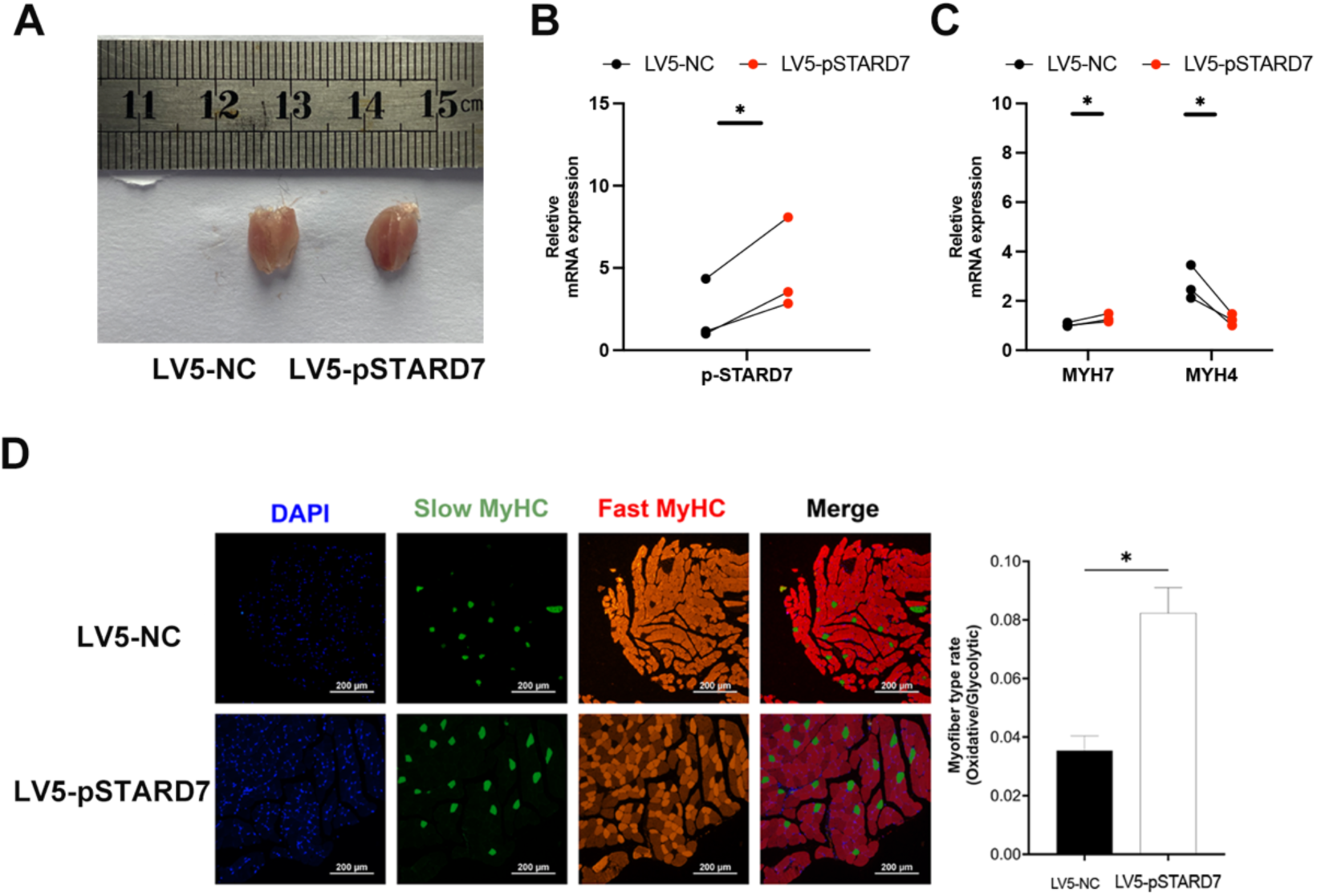
The monitor of fiber-type switching in vivo after the overexpression of pig STARD7 genes in mouse. (A) Left panel: Representative images of the GAS muscles from the left LV5-shNC or right LV5-pSTARD7 muscles. (B-C) qPCR showing gene expression changes. pSTARD7: pig STARD7 gene. (C) Representative image of immunofluorescence staining and statistical analysis of the proportion changes of fast-type fibers (green) and slow-type fibers (red).

**Fig. S10.**
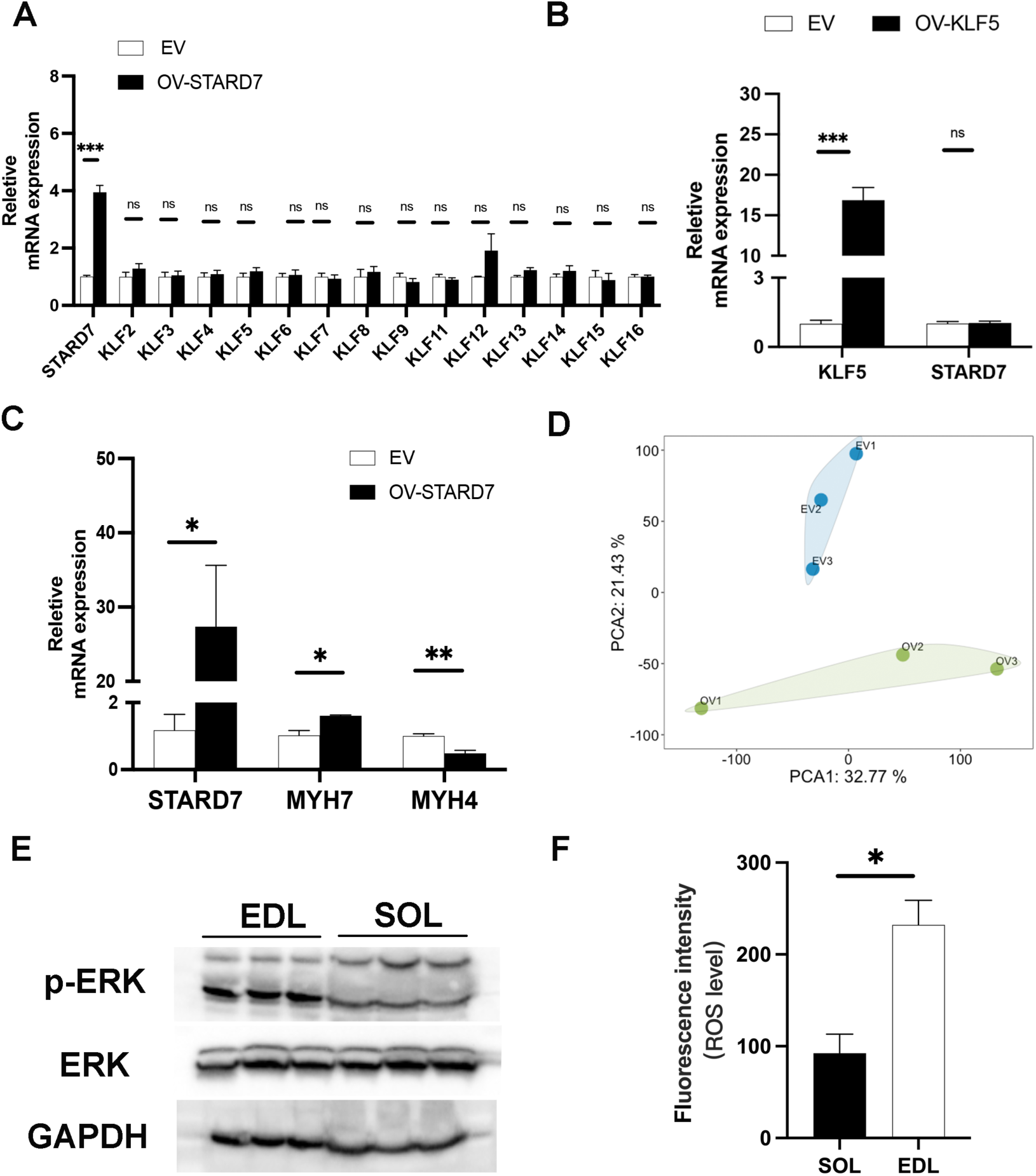
Transcriptome analysis of *STARD7* overexpression and quantitation of ROS level and ERK activity. (A) qPCR detecting changes in gene expression after *STARD7* overexpression. (B) qPCR detecting changes in gene expression after *KLF5* overexpression. (C) qPCR showing changes in gene expression following *STARD7* overexpression. (D) PCA plot of RNA-seq data from OV-*STARD7* and control PSCs. (E) WB results detecting the level of p-ERK and ERK in SOL and EDL muscles. (F) The florescence intensity of ROS in SOL and EDL. Excitation wavelength: 488nm, emission wavelength: 525nm.

